# Mitochondrial Ultrastructure Is Coupled to Synaptic Performance at Axonal Release Sites

**DOI:** 10.1101/216093

**Authors:** Csaba Cserép, Balázs Pósfai, Anett Dóra Schwarcz, Ádám Dénes

## Abstract

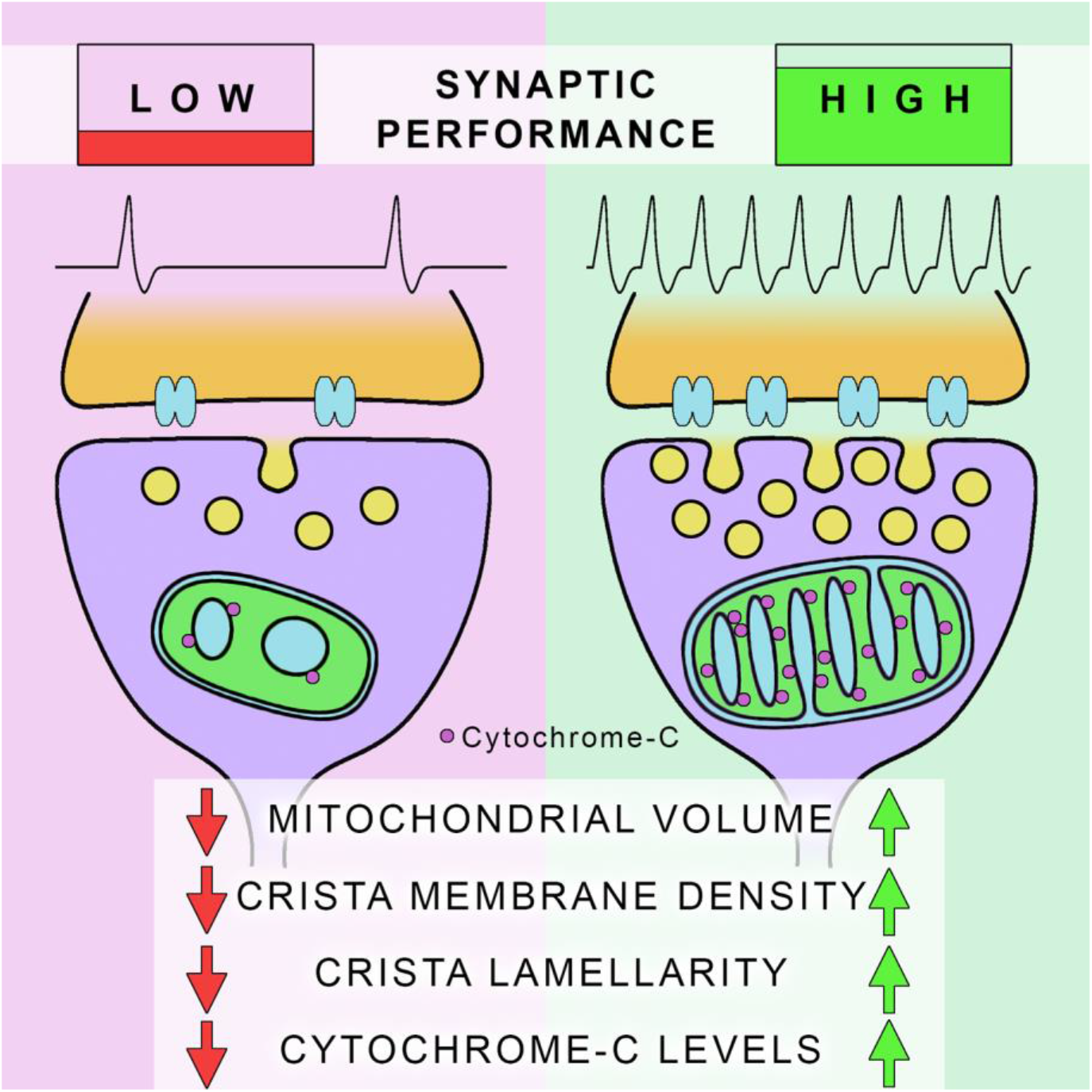

**SIGNIFICANCE STATEMENT:** Neuronal networks are highly dynamic structures: cellular morphology and synaptic strength is changing as information is processed and stored. This means that the demand for mitochondrial supply is changing in space and time in neurons. Since the architecture ultimately determines the output performance of these organelles, we hypothesized that the ultrastructure of axonal mitochondria could well be a major parameter of adjusting their performance to the actual energetic needs of the synapse. Our results – describing a cell-type independent and synapse-specific correlation between mitochondrial ultrastructure, mitochondrial molecular fingerprints and synaptic performance – highlight the importance of activity-dependent ultrastructural plasticity of these organelles in neurons, and suggest an even more prominent role for mitochondria in neuroplasticity, than it was thought previously.

**ABSTRACT:** Mitochondrial function in neurons is tightly linked with metabolic and signalling mechanisms that ultimately determine neuronal performance. The subcellular distribution of these organelles is dynamically regulated as they are directed to axonal release sites on demand, but whether mitochondrial internal ultrastructure and molecular properties would reflect the actual performance requirements in a synapse-specific manner, remains to be established. Here, we examined performance-determining ultrastructural features of presynaptic mitochondria in GABAergic and glutamatergic axons of mice and human. Using electron-tomography and super-resolution microscopy we found, that these features were coupled to synaptic strength: mitochondria in boutons with high synaptic activity exhibited an ultrastructure optimized for high rate metabolism and contained higher levels of the respiratory chain protein cytochrome-c than mitochondria in boutons with lower activity. The strong, cell-type independent correlation between mitochondrial ultrastructure, molecular fingerprints and synaptic performance suggests that changes in synaptic activity could trigger ultrastructural plasticity of presynaptic mitochondria, likely to adjust their performance to the actual metabolic demand.

## INTRODUCTION

About one-tenth of total grey matter volume is occupied by mitochondria, the powerhouses of cells (Toussaint and Kugler, 1989). These organelles form a rapidly changing, dynamic network in neurons (Barnhart, 2016). As primary energy generators, precise regulation of mitochondrial performance is fundamental for proper functioning of the central nervous system (CNS), due to the exceptionally high ATP-consumption of neuronal tissue. Mitochondria are also key regulators of intracellular calcium levels (Gunter et al., 2000; Hall et al., 2012) and contribute to signalling processes, the regulation of cell proliferation, migration, neuronal morphology and cell viability (Chandel, 2014). Since the main energy consumer in the CNS is synaptic transmission (Harris et al., 2012), intact mitochondrial ATP-production – together with calcium-regulation – is crucial for proper synaptic function and neuroplasticity (Billups and Forsythe, 2002; Gazit et al., 2016; Smith et al., 2016; Verstreken et al., 2005). Furthermore, the availability of presynaptic mitochondria at axonal release sites directly influences synaptic function (Rangaraju et al., 2014; Smith et al., 2016; Sun et al., 2013).

In highly dynamic neuronal networks cellular morphology and synaptic strength is changing in an activity dependent manner (Muller et al., 2002), resulting in altered neuronal demand for mitochondrial supply in space and time. Recent data also suggest that mitochondrial distribution within neurons is dynamically regulated to match the actual needs. *In vitro* studies confirmed, that motile mitochondria in dendrites as well as in axons are recruited and anchored to synapses whenever the local need for mitochondrial performance increases (Li et al., 2004; Sheng, 2014). It has also been shown in non-neuronal tissues that the inner ultrastructure of mitochondria determines their output performance. The surface area of crista-membrane (CM) in a given mitochondrial volume strongly correlates with mitochondrial oxygen consumption and cytochrome-oxidase activity (Else et al., 2004; Perkins et al., 2012, 2003; Sood et al., 2014). In addition to CM-density, the shape of cristae has also been shown to directly regulate mitochondrial performance output (Cogliati et al., 2013), and influence mitochondrial Ca^2+^-dynamics (Mannella et al., 2013).

Based on these data, we hypothesized that the ultrastructure of these organelles may be shaped to match the actual local needs in neurons. To test this idea with high-resolution imaging, we performed electron tomography and STORM super-resolution microscopy studies on mouse brain tissues, combined with serial-section transmission electron microscopy studies on postmortem human tissue samples. We examined presynaptic mitochondria in GABAergic and glutamatergic axons of the hippocampal formation. We found that CM-density, crista lamellarity and cytochrome-c density was higher in presynaptic mitochondria of the highly active fast-spiking parvalbumin-positive (PV) basket cells than in the mitochondria of the slow firing type-1 cannabinoid receptor-positive (CB1R) basket cells, which are the well-characterized archetypes of high-activity and low-activity interneurons, respectively. We also examined presynaptic mitochondria in the glutamatergic boutons of the perforant-path terminating zone in the dentate gyrus, and found a strong, cell-type independent correlation between mitochondrial volume, CM-density, crista lamellarity, cytochrome-c density and synaptic strength. The results from human post-mortem tissue also confirmed that this phenomenon is evolutionally conserved. Our data provide evidence for the logical assumption, that “stronger synapses use stronger mitochondria”, and point out the possibility of the activity-dependent ultrastructural plasticity of these organelles.

## MATERIALS AND METHODS

### Samples and tissue preparation

All experimental procedures were in accordance with the guidelines set by the European Communities Council Directive (86/609 EEC) and the Hungarian Act of Animal Care and Experimentation (1998; XXVIII, section 243/1998), approved by the Animal Care and Use Committee of the IEM HAS. 6 male C57BL/6 (RRID:IMSR_JAX:000664) mice and 4 CB1R knockout (CB1R-KO, Zimmer et al., 1999) mice (30-42 days old) were anesthetized with inhalation of isoflurane, followed by intraperitoneal injection of 0.05-0.1 ml of an anaesthetic mixture (containing 8.3mg/ml ketamine, 1.7 mg/ml xylazin-hydrochloride, 0.8 mg/ml promethazinium-chloride). Transcardial perfusions were performed in three different ways, depending on the type of experiment that followed. Animals used for immunofluorescent reactions (2 WT and 2 CB1R-KO mice for confocal scanning laser microscopy and STORM super-resolution microscopy) were perfused with 0.9% NaCl solution for 1 minute, followed by 4% freshly depolymerized paraformaldehyde (PFA) in 0.1 M phosphate buffer (PB) pH 7.4 for 40 minutes, and finally with 0.1 M PB for 10 minutes to wash the fixative out. For the immunogold reactions, we perfused other 2 WT and 2 CB1R-KO mice with saline for 1 minute, followed by 2% PFA and 1% glutar-aldehyde (GA) in 0.1 M Na-acetate buffer (pH=6) for 3 minutes, and then 2% PFA plus 1% GA in 0.1 M Borax-buffer (pH 8.5) for 40 minutes. In this case, the fixative was not washed out, but brains were removed, and postfixed overnight in the same fixative at 4 °C. For the experiments without immunostaining, we perfused 2 WT animals with saline for 1 minute, followed by 2% PFA and 2% GA in 0.1 M Na-acetate buffer (pH=6) for 3 minutes, and then 2% PFA plus 2% GA in 0.1 M Borax-buffer (pH 8.5) for 60 minutes. In this case again, the fixative was not washed out, but brains were removed and postfixed overnight in the same fixative at 4 °C. Blocks containing the dorsal hippocampi were dissected and coronal sections were prepared on a vibratome (VT1200S, Leica, Germany) at 20 μm thickness for STORM experiments, 50 μm thickness for immunofluorescent experiments, and 60 μm thickness for electron microscopy and electron tomography. Control human hippocampal tissue was obtained from one female (59-years-old) and one male (55-years-old) subject who died from causes not linked to brain diseases, and did not have a history of neurological disorders. The subjects were processed for autopsy in St. Borbála Hospital, Tatabánya, Department of Pathology. Informed consent was obtained for the use of brain tissue and for access to medical records for research purposes. Tissue was obtained and used in a manner compliant with the Declaration of Helsinki. All procedures were approved by the Regional and Institutional Committee of Science and Research Ethics of Scientific Council of Health (ETT TUKEB 31443/2011/EKU (518/PI/11)). Brains were removed 4-5 h after death. The internal carotid and the vertebral arteries were cannulated, and the brain was perfused first with physiological saline (using a volume of 1.5 L in 30 min) containing heparin (5 ml), followed by a fixative solution containing 4% paraformaldehyde, 0.05% glutaraldehyde and 0.2% picric acid (vol/vol) in 0.1 M PB, pH 7.4 (4–5 L in 1.5–2 h). The hippocampus was removed from the brain after perfusion, and was postfixed overnight in the same fixative solution, except for glutaraldehyde, which was excluded. Blocks were dissected, and 60 μm thick sections were prepared on a vibratome (VT1200S, Leica, Germany).

### Primary antibodies

To detect CB1Rs, we used either a goat polyclonal antibody or a rabbit polyclonal antibody (gifts from Prof. Ken Mackie). For the staining of parvalbumin (PV), we used a rabbit polyclonal antibody (PV27, Swant, RRID:AB_2631173). The specificity of this antibody was tested on PV-KO tissue (Swant). To label GABAergic boutons, we used a KO-verified guinea-pig anti-vesicular GABA Transporter (vGAT) antibody (131004, Synaptic Systems, RRID:AB_887873). To visualize cytochrome-c (CytC), we used a mouse monoclonal antibody (clone 6H2.B4, 612302, Biolegend, RRID:AB_315775). The specificity of this antibody has been previously tested and described (Gulyás et al., 2006). To detect TOM20, we used a rabbit polyclonal antibody (Santa Cruz Biotechnology, cat. no. s-11415, RRID:AB_2207533, Barna et al., 2016). To label glutamatergic presynaptic boutons, we used a polyclonal guinea-pig antibody, raised against a synthetic peptide from rat vGluT1 (VG1), which peptide does not overlap with the sequence of vGluT2; (Millipore, AB5905, RRID:AB_2301751). The specificity of this antibody has also been described, and it gave identical staining with a well-characterized rabbit anti-VG1 antibody (Melone et al., 2005). A rabbit polyclonal antibody (Synaptic Systems, 160003, RRID:AB_887730), raised against the 1-186 amino-acid residues of Homer1 was used to label the postsynaptic density of glutamatergic synapses. This antibody stains selectively the postsynaptic density of glutamatergic synapses (Andreska et al., 2014).

All antibodies gave the previously described and thus expected staining patterns for their respective epitopes. For further control of our stainings, we extensively tested the possible crossreactivity of the fluorescent secondary antibodies used in multiple labeling experiments. No cross-reactivity was found in any of the cases. Selective labeling, resembling that obtained with the specific antibodies, could not be detected if primary antibodies were omitted.

### Immunofluorescent labeling and confocal laser-scanning microscopy

Before the immunofluorescent staining, the 50 μm thick sections were washed in PB and Tris-buffered saline (TBS, 0,05 M Tris, 0,9% NaCl, pH=7.4). This was followed by blocking for 1 hour in 1% human serum albumin (HSA) and 0.1% Triton X-100 dissolved in TBS. After this, sections were incubated in mixtures of primary antibodies 1.) rabbit and goat anti-CB1R, both diluted in TBS 1:500; or 2.) rabbit anti-parvalbumin, 1:2000; goat anti-CB1R, 1:500; and guinea-pig anti-vGAT, 1:2000; overnight at room temperature. After incubation, sections were washed in TBS, and sections were incubated overnight at 4 °C in the mixture of 1.) Alexa488 conjugated donkey-anti-rabbit, (1:500, Life Technologies) and Alexa594 conjugated donkey-anti-goat (1:500, Invitrogen) antibodies; or 2.) Alexa488 conjugated donkey-anti-rabbit, (1:500, Life Technologies), Alexa594 conjugated donkey-anti-goat (1:500, Invitrogen) and Alexa647 conjugated donkey-anti-guinea-pig, (1:500, Jackson Immuno Research), all diluted in TBS. Secondary antibody incubation was followed by washes in TBS, PB, and the sections were treated with DAPI for 1.5 hours (1:10000, in PB). Finally, we washed the sections, mounted them onto glass slides, and coverslipped with Aqua-Poly/Mount (Polysciences). Immunofluorescence was analyzed using a Nikon Eclipse Ti-E inverted microscope (Nikon Instruments Europe B.V., Amsterdam, The Netherlands), with a CFI Plan Apochromat VC 60XH oil immersion objective (numerical aperture: 1.4) and an A1R laser confocal system. We used 405, 488, 561 and 642 nm lasers (CVI Melles Griot), and scanning was done in line serial mode. Image stacks were obtained with NIS-Elements AR software, and deconvolved using Huygens Professional software (www.svi.nl).

### STORM super-resolution microscopy

Before the immunofluorescent staining for the super-resolution experiments, the 20 μm thick sections were washed in PB and TBS. This was followed by blocking for 1 hour in 1% HSA dissolved in TBS. After this, sections were incubated in mixtures of primary antibodies 1.) goat anti-CB1R, 1:500; rabbit anti-PV, 1:2000; and mouse anti-CytC, 1:3000; 2.) rabbit anti-TOM20, 1:500 and mouse anti-CytC, 1:3000; or 3.) guinea-pig anti-VG1, 1:5000; rabbit anti-Homer1, 1:2000 and mouse anti-CytC, 1:3000; overnight at room temperature. After incubation, sections were washed in TBS, and sections were incubated overnight at 4 °C in the mixtures of secondary antibodies 1.) DyLight405 conjugated donkey-anti-goat, (1:500, Jackson), Alexa488 conjugated donkey-anti-rabbit, (1:500, Life Technologies), and Alexa647 conjugated donkey-anti-mouse, (1:500, Jackson); CF568 conjugated donkey-anti-rabbit (1:500, Jackson) and A647 conjugated donkey-anti-mouse (1:500, Jackson); or 3.) DyLight405 conjugated donkey-anti-guinea-pig, (1:500, Jackson), Alexa488 conjugated donkey-anti-rabbit, (1:500, Life Technologies) and Alexa647 conjugated donkey-anti-mouse, (1:500, Jackson). Secondary antibody incubation was followed by washes in TBS, PB, and hippocampi were cut out with scalpels under buffer, sections were dried on clean coverslips, and stored for maximum 3 weeks at room temperature in a dry environment before imaging. Imaging medium was mixed from 80 μl DPBS, 10 μl MEA solution, 10 μl 50% glucose solution, and 1 μl GLOX solution. Coverslips with the dried sections were mounted onto microscope slides with 25 μl of freshly prepared imaging medium, and sealed with nail-polish. 3D direct-STORM (dSTORM) acquisition protocol was used as described before (Barna et al., 2016). The imaging setup was built around a Nikon Ti-E inverted microscope equipped with a Perfect Focus System, with an Andor iXon Ultra 897 EMCCD camera, a C2 confocal head, 405, and 488 nm lasers (Melles Griot 56RCS/S2780, Coherent Sapphire), and a high power 647 nm laser (300 mW, MPB Communications VFL-P-300-647). A high NA 100x oil immersion objective (Nikon CFI SR Apochromat TIRF 100x oil, 1.49 NA) was used for imaging. Emitted light was let through a 670/760 nm bandpass filter to reach the detector. We used the NIS-Elements AR 4.3 with N-STORM 3.4 software for acquisition and analysis. After selecting the region of interest, confocal image stacks were acquired containing 10 focal planes with 80 x 80 x 150 nm voxel size in x, y and z, respectively. This was followed by bleaching the fluorophores in the STORM channels (561 and 647) with high intensity laser illumination, and running dSTORM acquisition using oblique (near-TIRF) illumination. Acquisition in the two channels was done in sequential mode. Confocal stacks were deconvolved (Huygens Professional), and – together with the STORM molecule lists – processed for further analysis in VividSTORM software (see „Analysis”). Imaging was performed with identical parameters (depth in the section, laser intensities, etc.) for all samples.

### Pre-embedding immunoelectron-microscopy

For the immunogold-labeling of CB1Rs, sections were washed in PB, treated with 0.5% sodium-borohydride for 13 minutes, and further washed in PB and TBS. After this, we incubated the sections in 1% HSA diluted in TBS. Then the sections were incubated for 48 hours in rabbit anti-CB1R primary antibody diluted in TBS (1:500). After repeated washes in TBS, sections were treated with blocking solution (Gel-BS) containing 0.5% cold water fish skin gelatin (Aurion, Netherlands) and 0.5% HSA in TBS for 1 h. This was followed by 24 h incubation in 1.4 nm gold conjugated goat anti-rabbit Fab-fragment (1:200, NanoProbes) diluted in Gel-BS. After intensive washes in TBS and 0.1 M PB sections were treated with 2% glutar-aldehyde in 0.1 M PB for 15 minutes to fix the gold particles into the tissue. This was followed by further washes in 0.1 M PB and enhancement conditioning solution (ECS, Aurion), gold particles were intensified using the silver enhancement solution (SE-EM, Aurion) for 40-60 minutes at room temperature. After subsequent washes, sections were treated with 0.5 % osmium-tetroxide in PB for 20 minutes on ice. Then sections were dehydrated in ascending ethanol series and acetonitrile, and embedded in epoxy resin (Durcupan, ACM, Fluka, Buchs, Switzerland). During dehydration sections were treated with 1% uranyl-acetate in 70% ethanol for 20 minutes. After polymerization, 60 or 200 nm thick sections were cut using a Leica EM UC6 ultramicrotome (Nussloch, Germany), and picked up on formvar-coated single-slot copper grids. 60 nm thick sections were examined using a Hitachi H-7100 electron microscope (Tokyo, Japan) and a side-mounted Veleta CCD camera (Olympus Soft Imaging Solutions), while 200 nm thick sections were used for electron tomographic examinations.

### Serial-section electron-microscopy of human samples

After washing the 60 μm thick sections in 0.1 M PB, sections were treated with 0.5 % osmium-tetroxide in PB for 20 minutes on ice. Then sections were dehydrated in ascending ethanol series and acetonitrile, and embedded in epoxy resin (Durcupan, ACM, Fluka, Buchs, Switzerland). During dehydration sections were treated with 1% uranyl-acetate in 70% ethanol for 20 minutes. After polymerization, 60 or 200 nm thick sections were cut using a Leica EM UC6 ultramicrotome (Nussloch, Germany), and picked up on formvar-coated single-slot copper grids. 60 nm thick sections were examined using a Hitachi H-7100 electron microscope (Tokyo, Japan) and a side-mounted Veleta CCD camera (Olympus Soft Imaging Solutions). Serial sections were examined, and mitochondria-containing glutamatergic boutons establishing asymmetric synapses were collected in the CA1 area, and reconstructed from long, serial sections. Measurement of synaptic areas and mitochondrial volumes were performed with Fiji/ImageJ software.

### Electron tomography

For the electron tomographic investigation, we used either 200 nm thick sections from the hippocampal CA1 region from the anti-CB1R immunogold stained material (see: “Preembedding immunoelectron-microscopy”), or 100 nm thick sections of unstained tissue from the dentate gyrus. In the latter case the sections were washed thoroughly in PB after postfixation (see “Samples and tissue preparation”), and treated with 0.5 *%* osmium-tetroxide in PB for 20 minutes on ice. Then sections were dehydrated in ascending ethanol series and acetonitrile, and embedded in epoxy resin (Durcupan). During dehydration sections were treated with 1% uranyl-acetate in 70% ethanol for 20 minutes. Before electron tomography, serial sections on single-slot copper grids were photographed with a Hitachi H-7100 electron microscope and a Veleta CCD camera. Serial sections were examined at lower magnification, and mitochondria-containing perisomatic synaptic boutons from the CA1 area, or mitochondria-containing glutamatergic synaptic boutons from the outer two-thirds of DG molecular layer were selected. At this magnification inner mitochondrial ultrastructure could not be observed on the TEM images from 100 and 200 nm thick sections, securing the unbiasedness of sampling. In the CA1 experiments, CB1R immunogold staining was examined at higher magnification, and boutons were divided into CB1R+ and CB1R-(PV) categories. In the DG experiments, the boutons were reconstructed from the serial TEM image series, synapse sizes and mitochondrial volumes were measured. For each bouton, the section containing the largest mitochondrial cross section was chosen for electron tomography. After this, grids were put on drops of 10% HSA in TBS for 10 minutes, dipped in distilled water (DW), put on drops of 10 nm gold conjugated Protein-A in DW (1:3), and washed in DW. Finally, we deposited 5-5 nm thick layers of carbon on both sides of the grids. Electron tomography was performed using a Tecnai T12 BioTwin electron microscope equipped with a computer-controlled precision stage (CompuStage, FEI). Acquisition was controlled via the Xplore3D software (FEI). Regions of interest were preilluminated for 6 minutes to minimize further shrinkage. Dual-axis tilt series were collected at 2-degree increment steps between -65 and +65 degrees at 120 kV acceleration voltage and 23000x magnification with -1.6 – -2 μm objective lens defocus. Reconstruction was performed using the weighted backprojection algorithm in the IMOD software package (RRID:SCR_003297, Kremer et al., 1996). Isotropic voxel size was 0.49 nm in the reconstructed volumes. After combining the reconstructed tomograms from the two axes, the nonlinear anisotropic diffusion (NAD) filtering algorithm was applied to the volumes. Segmentation of mitochondrial volumes has been performed on the virtual sections using the 3Dmod software, and measurements were done on the scaled 3D models.

### Methodological considerations

Electron tomography has become the primary method for studying biological structures in their native environment with the highest resolution. For these type of studies, it is critical to use tissue preparation techniques that most reliably preserve the physiological state. Throughout the last decades, transcardial perfusion with aldehydes has become the gold standard method of electron microscopic tissue preparation when cryo-fixation is not an option (as in the case of intact brains). However, this technique is known to introduce some structural artefacts (decreased extracellular volume, decreased number of docked synaptic vesicles) into the sample when compared with rapid cryo-fixation (Korogod et al., 2015). Because of this, one needs to consider whether their results and the interpretation of these results are affected by these differences, or not.

In our case, there are two main reasons for being convinced, that our results are unaffected by this problem. Firstly, the parameters we set out to explore – namely the surface area and architecture of mitochondrial crista-membranes – have been described to be unaffected by perfusion and aldehyde fixation when compared with rapid cryo-fixation and cryo-electron tomography (Nicastro et al., 2000; Perkins et al., 1998). Secondly, we always made comparisons within one system, and compared mitochondria that were sampled equally for each group from the same tissue volumes, meaning that they must have been affected identically, if anyhow. These arguments validate our approach, and enable us to draw biologically relevant conclusions from our data.

### Controls

To validate the specificity and sensitivity of our labeling approach, we performed immunofluorescent reactions with two different CB1R antibodies in wild-type (WT) and CB1R-knockout (KO) mice (Fig. 1-1. A, B). The two antibodies showed completely overlapping staining in the WT animals, but were totally absent in the KO animals (n=2 WT and 2 KO mice). For the electron tomographic examinations, we performed immunogold labeling against CB1R (Fig. 1. B), and quantified labeling density in WT and CB1R-KO animals (Fig. 1. C). It was found to be 4.87 particle/μm on presynaptic CB1R+ boutons in WT, and 0.00 particle/μm in KO animals (tested along 38.6 μm membrane in 2 WT, and tested along 74.8 μm presynaptic bouton membrane in 2 KO animals). This would correspond to an average of 12 gold particles on a single electron microscopic (EM) section of a CB1R+ bouton in WT animals, reflecting the exceptionally high specificity and sensitivity of CB1R labeling, and confirming that this method can reliably distinguish between the two perisomatic bouton populations on serial EM images. To estimate the necessary pre-irradiaton of plastic embedded sections before electron tomography, we measured the dynamics of tissue shrinkage on 100 and 200 nm thick sections. Shrinking happened mainly in the first 2 minutes of sample irradiation, and decreased to a negligible rate after 4-5 minutes (Fig. 1-1. C). After these measurements, we set a uniform preirradiation time of 6 minutes in our experiments.

**Figure 1.**
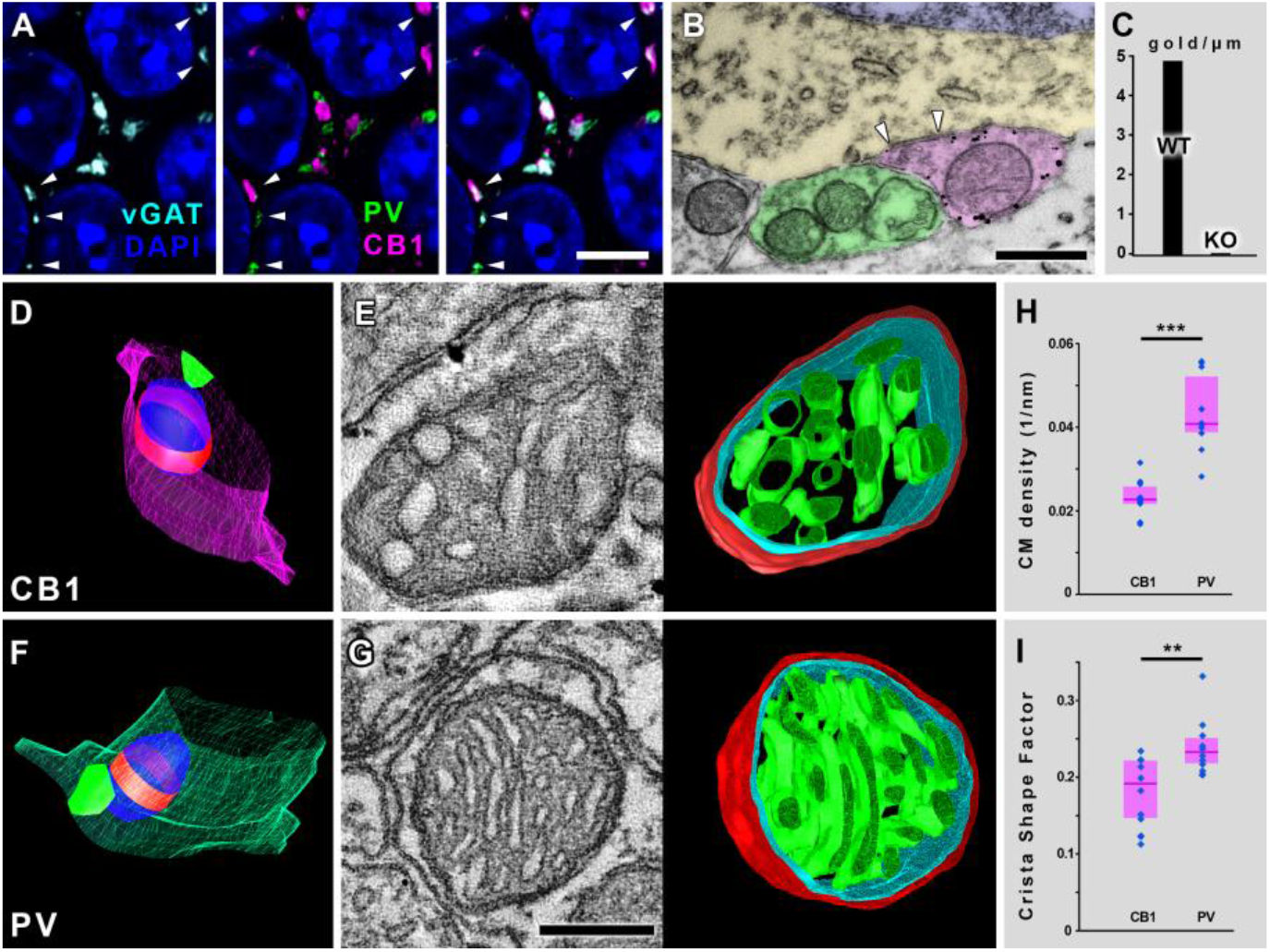
Electron tomography reveals robust ultrastructural differences between axonal mitochondria of fast-spiking and regular-spiking basket cells. (A) Confocal images show that perisomatic synaptic boutons are either PV- or CB 1R-positive in the hippocampal CA1 region. Pyramidal cell nuclei are labeled with DAPI (blue), vGAT-immuno-reactive puncta represent GABAergic vesicle pools (cyan), PV-labeling is green and CB 1R-labeling is magenta. (B) Transmission electron micrograph shows that CB 1R-immunogold labeling reliably differentiates the two perisomatic bouton populations. Black granules are the silver-intensified CB1R-immunogold particles, white arrowheads mark a synapse. Pseudocolors: CB1R+ bouton – magenta, PV bouton – green, pyramidal cell cytoplasm – yellow and nucleus – blue. (C) CB1R-immunogold labeling was absent in CB1R-KO animals (tested membrane length: 38 μm in two WT, and 74.8 μm in two CB1R-KO mice). (D, F) Representative 3D-models of serial EM reconstructed segments from a synaptic CB1R+ bouton (D), and a synaptic PV bouton. Bouton membrane is magenta for CB1R-bouton, green for PV-bouton, synapses are vivid-green, mitochondria are blue, red belts represent those sections of the organelles that were reconstructed through electron tomography. (E) 1.5 nm thick electron tomographic section and 3D-model of the reconstructed mitochondrion from the bouton depicted in D (mitochondrial outer membrane is red, inner boundary membrane is cyan, crista membrane is green). (G) 1.5 nm thick electron tomographic section and 3D-model of the reconstructed mitochondrion from the bouton depicted in F. (H) Crista-membrane density is significantly higher in mitochondria of PV-boutons than in those from CB1R+ boutons (Mann-Whitney U-test, p=0.0002, n=20 mitochondria from 2 mice). (I) Cristae are significantly more lamellar in mitochondria of PV-boutons than in those from CB1R+ boutons, as verified by the higher crista shape factor values (Mann-Whitney U-test, p=0.0090, n=20 mitochondria from 2 mice). (H, I) Blue dots represent values from individual mitochondria, magenta rectangles represent interquartile ranges, deep-magenta lines mark median values. Scale bars are 6 μm in A, 500 nm in B and 100 nm in G and for E. The 3D models on E and G are not displayed on the same scale. See also Figures 1-1 and 1-2.

### Analysis and statistics

In the immunogold experiments the gold particles were considered to be membrane-associated, if they were not further away from the membrane than 40 nm, because the epitope (the C-terminus of the CB1R) is located a few nanometers intracellularly, and the length of the primary and secondary IgG antibody molecules are ~15 nm each.

During segmentation of tomographic volumes, z-scaling was calculated from the thickness difference of the reconstructed volume and the original section thickness, and applied to the 3D models to compensate for z-shrinkage. Mesh surface areas and volumes inside meshed objects were measured with the “imodinfo” program. To measure synaptic area in the serial TEM images, the length between the edges of anatomically defined synapses was measured on subsequent images of the sampled synapses, and the sum of the measured lengths from each synapse was multiplied by the section thickness.

In the confocal triple labeling experiments to visualize VG1-Homer-CytC signals, segmentation was performed separately for each channel to ensure unbiasedness.

STORM analysis was performed in the VividSTORM open source software (Barna et al., 2016). Localisation points exceeding a given photon count were counted as specific SLPs. Molecule lists were exported from NIS in txt format, and the 3 image planes of the ics-ids file pairs from the deconvolved confocal stacks matching the STORM-volume were converted to the ome-tiff format using Fiji software. Confocal and corresponding STORM images were fitted in VividSTORM. Convex 2D hulls were automatically constructed around individual (not in fission or in fusion) mitochondria in both channels (568-TOM20 and 647-CytC), and organelle areas were correlated. During the examination of mitochondria in perisomatic boutons, the identity (CB1R+ vs. PV+) of the boutons with mitochondria was determined in the confocal channels, and a convex 2D hull was constructed around the corresponding SLP cluster. The number of CytC SLPs was divided by the area of this convex hull to calculate the SLP-density within mitochondria. During the examination of mitochondria in perforant pathway boutons, VG1+ boutons were identified with presynaptic mitochondria, and the size of the corresponding active zone was determined by 3D-reconstruction of the Homer confocal signal. CytC SLP-density was determined as described above.

When data populations did not have a Gaussian distribution according to the Shapiro-Wilks W test, we reported non-parametric statistical features (median, interquartile range), compared two independent groups with the Mann-Whitney U-test, and Spearman correlation coefficients were calculated. When data populations showed normal distribution, correlations were assessed using linear regression fit to the data (R indicates Pearson’s correlation coefficient). Statistical analysis was performed with the Statistica 13.1 package (RRID:SCR_014213, Dell), differences with p<0.05 were considered significant throughout this study. Since data from different animals belonging to the same group were not statistically different in any case, they were pooled.

## RESULTS

### Electron tomography reveals robust ultrastructural differences between presynaptic mitochondria of fast-spiking and regular-spiking GABAergic neurons

In order to test whether any correlation exists between synaptic performance and mitochondrial ultrastructure, we defined a “low performance” and a “high performance” group within hippocampal GABAergic as well as glutamatergic synaptic populations, based on the level of their activity. In the hippocampal CA1 area, the perisomatic GABAergic inputs of principal cells can be divided into two, morphologically and functionally well-characterized populations: the CB1R-positive, regular-spiking interneurons show low level of synaptic activity, whereas the PV-positive, fast-spiking interneurons are capable of performing exceptionally high synaptic activity. In fact, these GABAergic populations are the two well-known archetypes of the slow/modulatory, and the highly energized/fast “clockwork” type interneurons, respectively (Gulyás et al., 2006; Kann et al., 2014; Klausberger et al., 2005). The average *in vivo* firing frequency of the CB1R+ interneurons in the hippocampus is 2-10 Hz, whereas the PV+ interneurons can fire bursts up to 150 Hz during ripples (Klausberger et al., 2005; Lapray et al., 2012), making these two populations ideal candidates for “low performance” (LP) and “high performance” (HP) GABAergic boutons, respectively. Since the perisomatic synaptic boutons contacting pyramidal cells in the hippocampal CA1 region originate exclusively (99%) from these two cell types (Takács et al., 2015), labeling one of them is sufficient to reliably identify both types. To verify the exclusive and non-overlapping contribution of the two populations with our antibodies as well, we performed immunofluorescent labeling of GABAergic varicosities for vesicular GABA transporter (vGAT), the two bouton-populations for CB1R and PV, and cell nuclei with DAPI (Fig. 1., A). We found, that 54.5% of all GABAergic boutons on pyramidal cell somata were PV+, 45% CB1R+ and only 0.5 % were double-negative (n=200 boutons in 2 mice), which numbers are in a very good agreement with previous data (Takács et al., 2015). For the electron tomographic examinations, we performed immunogold labeling against CB1R (Fig. 1. B). Once the mitochondrion-containing boutons from both populations were sampled, the presence of a synapse was confirmed for each bouton (Fig. 1. D, F). The sections containing the largest crosssections of mitochondria were processed for dual-axis electron tomography and reconstruction, which revealed robust ultrastructural differences between these organelles, depending on which population they belonged to (Fig. 1. E, G, Fig 1-2.). Crista-membrane density (surface area of crista membrane/mitochondrial volume) was significantly higher (by 79.7%) in mitochondria of PV boutons (median: 0.0408 nm^-1^, interquartile range: 0.0389-0.0519) than in those from CB1R+ boutons (median: 0.0227 nm^-1^, interquartile range: 0.0218-0.0256, Mann-Whitney U-test, p=0.0002, n=20 mitochondria from 2 mice, Fig. 1. H). When we multiplied these crista membrane density values with the median volume of presynaptic mitochondria (119x10^6^ nm^3^ in PV+ and 73.4 x10^6^ nm^3^ in CB1R+ boutons from Takács et al., 2015) we found a striking difference, showing that the average PV+ bouton possesses 2.9 times more mitochondrial crista membrane surface than the average CB1R+ bouton. The organisation of the cristae also differed, since these structures were significantly more lamellar in mitochondria of PV-boutons than in those from CB1R+ boutons, as verified by the 22%-higher crista shape factor values (surface area of crista membrane/crista lumen volume, referred to as lamellarity throughout the paper), PV median: 0.233 nm^-1^, interquartile range: 0.218-0.251, CB1R median: 0.191 nm^-1^, interquartile range: 0.147-0.220, Mann-Whitney U-test, p=0.0090, n=20 mitochondria from 2 mice, Fig. 1. I). These results suggest a possible correlation between the ultrastructure of presynaptic mitochondria and the performance of the synapses.

### STORM super-resolution microscopy confirms higher mitochondrial cytochrome-c density in boutons of fast-spiking basket cells than in regular-spiking ones

To assess whether the observed ultrastructural differences are accompanied by any changes in respiratory chain protein expression, we turned to a combined stochastic optical reconstruction microscopy (STORM) and confocal laser-scanning microscopy (CLSM) method, and performed precise quantitative assessment of cytochrome-c (CytC) levels in mitochondria. This protein is an indispensable element of the electron-transport chain, as it carries electrons to the cytochrome-c oxidase enzyme. CytC expression directly controls oxidative phosphorylation (Wilson et al., 2014), and its levels are indicative of mitochondrial performance and show good correlation with neuronal activity (Gulyás et al., 2006; Kann et al., 2014). To take advantage of the unprecedented 20-30 nm lateral localisation precision and qunatitative nature of the STORM super-resolution technique (Huang et al., 2008) and delineate mitochondrial borders with high fidelity, we performed double labeling experiments. If the distribution of CytC is homogenous within individual mitochondria, then the CytC superresolution localisation point (SLP) clusters would designate the borders of these organelles in a reliable way. In this case, one could measure the sizes of these clusters and compare the SLP-density values between different mitochondria. To test this, we stained for TOM20 (central component of the translocase of outer membrane receptor complex) and CytC (Fig. 2. A). The measured sizes of individual mitochondria in the two STORM channels showed a very strong correlation (R=0.98, n=48 mitochondria from 2 mice, Fig. 2. B), excluding the non-homogenous sub-organelle distribution of CytC labeling. The median number of CytC SLPs labeling a single mitochondrion was 217 (interquartile range: 97-330). These results confirmed, that the CytC-labeling and STORM imaging can be used to delineate the borders of mitochondria in a precise and reliable way. Next, we performed a multi-color immunofluorescent labeling for PV, CB1R and CytC, and imaged the samples using correlated confocal and STORM super-resolution microscopy (Fig. 2. C-G). The CytC-labeling was recorded with the STORM system as well. The STORM SLP-s were overlayed the confocal images with the VividSTORM software (Fig. 2. D, F). The analysis showed, that mitochondria in the PV+ boutons were larger than those in the CB1R+ boutons, as the 2D areas of the convex hulls – fitted around SLP-clusters – were 75.4% larger in the former group (median: 0.287 μm^2^, interquartile range: 0.158-0.368) than in the latter (median: 0.164 μm^2^, interquartile range: 0.098-0.203, Mann-Whitney U-test, p=0.0085, n=57 mitochondria from 2 mice, Fig. 2. H). More importantly we found, that mitochondria in the PV+ boutons contained CytC SLPs in a 22.7% higher density (median: 565.7 SLP/μm^2^, interquartile range: 434.8-725.7) than those in the CB1R+ boutons (median: 461.1 SLP/μm^2^, interquartile range: 378.8-612.0, Mann-Whitney U-test, p=0.0384, n=57 mitochondria from 2 mice, Fig. 2. I). These results confirm that the internal structural differences between mitochondria in the two bouton populations are accompanied by quantitative differences in molecular fingerprints as reflected by the levels of the respiratory chain protein cytochrome-c.

**Figure 2.**
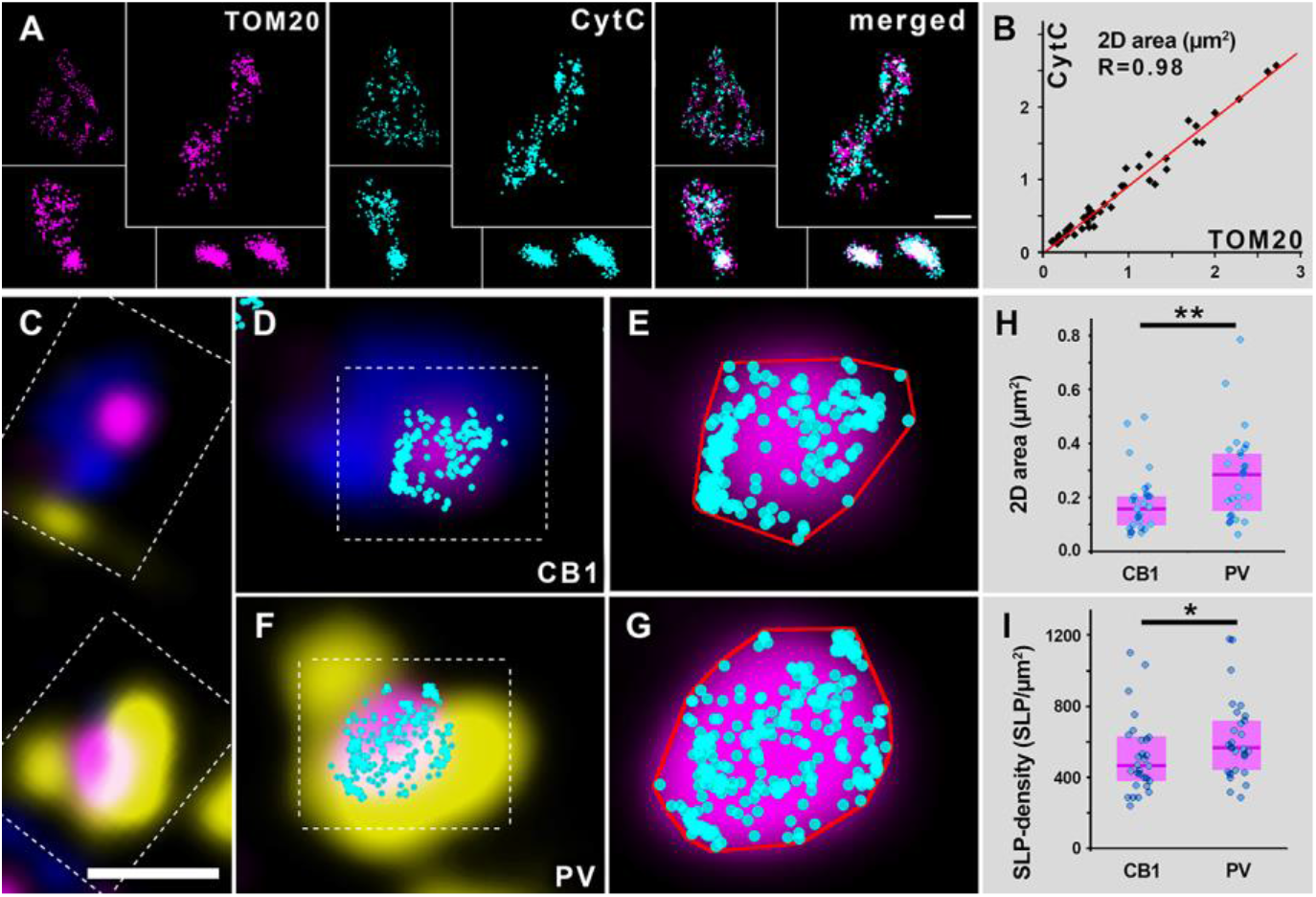
Axonal mitochondria of fast-spiking basket cells express higher levels of cyto-chrome-c than those of regular-spiking basket cells’. (A) STORM superresolution imaging confirms a near-complete overlap between TOM20 and cytochrome-c labeled areas. (B) Scatterplot shows a strong correlation (R=0.98, n=48 mitochondria from 2 mice) between the measured areas in the two channels of individual mitochondria. Each dot corresponds to a single mitochondrion. (C) Confocal image shows a CB1R+ (blue) and a PV+ (yellow) perisomatic bouton, both containing a mitochondrion, labeled for cytochrome-c (magenta). (D) CB1R+ bouton enlarged from C with overlaid STORM cytochrome-c localisation points (cyan). (E) Confocal and STORM image of mitochondrion enlarged from D. (F) PV+ bouton enlarged from C with overlaid STORM cytochrome-c localisation points. (G) Confocal and STORM image of mitochondrion enlarged from D. Red lines in E and G mark the 2D convex hulls generated around the localisation points. (H) Mitochondria in the PV+ boutons are larger than those in the CB1R+ boutons, as the 2D areas of the convex hulls are significantly larger in the former group (Mann-Whitney U-test, p=0.0085, n=57 mitochondria from 2 mice). (I) Mitochondria in the PV+ boutons contain cytochrome-c in a higher density than those in the CB1R+ boutons (Mann-Whitney U-test, p=0.0384, n=57 mitochondria from 2 mice). Blue dots represent values from individual mitochondria, magenta rectangles represent interquartile ranges, deep-magenta lines mark median values. Scale bar in A is 800 nm for upper-left mitochondrion, 400 for all others; scale bar in C is 1 μm, 600 nm for D and F and 170 nm for E and G.

### Ultrastructural parameters of presynaptic mitochondria in glutamatergic boutons are coupled to synaptic strength

To confirm that the observed correlations between molecular and structural mitochondrial properties and synaptic features are independent of the cell types examined, we investigated glutamatergic synaptic boutons in the dentate gyrus. One of the best characterized structure-function correlations in neuroscience is that synaptic strength is strictly and faithfully linked to morphological/ultrastructural parameters (Bourne and Harris, 2012; Buchs and Muller, 1996; Cheetham et al., 2014; Desmond and Levy, 1988; Holtmaat and Svoboda, 2009). The release probability of synapses scales linearly with the size of the active zone (Holderith et al., 2012), and long-term potentiation leads to an increase in the size of synaptic active zone, the number of perforated synapses and also in the number of boutons with multiple active zones (Geinisman, 2000; Popov et al., 2004). Based on this fundamental morpho-functional correlation, the strength of glutamatergic synapses can be determined purely by morphological criteria in a very reliable way, enabling us to define a “low performance” and a “high performance” group within a specific glutamatergic synaptic bouton population. We randomly collected samples from the outer two-thirds of the dentate gyrus molecular layer, where the vast majority of glutamatergic contacts are made by perforant path axons (Amaral et al., 2007), and reconstructed synaptic boutons from serial EM sections together with the postsynaptic neuronal processes (Fig. 3. A, C). The median active zone area was found to be 0.070 μm^2^ (0.042-0.123 μm^2^ interquartile range, n=35 synaptic boutons in 2 mice, Fig. 3-1. A). About half of these boutons contained mitochondria, and the active zone area in this group did not differ significantly from the previous group, as it was found to be 0.093 μm^2^ (0.048-0.171 μm^2^ interquartile range, Mann-Whitney U-test, p=0.267; n=19 synaptic boutons in 2 mice, Fig. 3-1. A). Based on these data and the strict correlation between synaptic ultrastructure and performance, we grouped the reconstructed mitochondria-containing varicosities as follows: boutons were considered to be “low performance” (LP), if they had a single, non-perforated active zone with an area less than 0.09 μm^2^, and boutons were considered to be „high performance” (HP), if they had multiple and/or peforated active zone(s) with an area larger than 0.09 μm^2^ (Fig. 3-1. A). We measured the mitochondrial volumes in the 3D-models, and found that presynaptic mitochondria in HP-boutons were significantly larger than those in LP-boutons (HP-median: 41.8x10^6^ nm^3^, 25x10^6^-46.5x10^6^ interquartile range, LP-median: 18.9x10^6^ nm^3^, 13x10^6^-27.6x10^6^ interquartile range, 121.6% difference, Mann-Whitney U-test, p=0.0199, n=19 boutons from two mice; Fig. 3. E). The sections containing the largest cross-sections of presynaptic mitochondria of these boutons were processed for dual-axis electron tomography and reconstruction, which revealed robust ultrastructural differences between these organelles, depending on which population they belonged to (Fig. 3. B, D, Fig. 3-2.). Crista-membrane density (surface area of crista membrane/mitochondrial volume) was significantly higher (by 110.1%) in mitochondria of HP-boutons (median: 0.0552 nm^-1^, interquartile range: 0.05090–0587) than in those from LP-boutons (median: 0.0262 nm^-1^, interquartile range: 0.0258–0.0290, Mann-Whitney U-test, p=0.0009, n=19 mitochondria from 2 mice, Fig. 3. F). We then multiplied these crista membrane density values with the median volume of presynaptic mitochondria (41.8x10^6^ nm^3^ in HP and 18.9x10^6^ nm^3^ in LP boutons), and found a striking difference, showing that the average HP bouton possesses 4.67 times more mitochondrial crista membrane surface than the average LP bouton. The organisation of the cristae also differed significantly, since these structures were more lamellar in mitochondria of HP-boutons than in those from LP-boutons, as verified by the 37.8% higher crista shape factor values (surface area of crista membrane/crista lumen volume) in the former group (median: 0.268, interquartile range: 0.237–0.301), than in the latter (median: 0.195, interquartile range: 0.190–0.219, Mann-Whitney U-test, p=0.0080, n=19 mitochondria from 2 mice, Fig. 3. G). To verify the robustness of our results, we pooled data from HP and LP boutons, and examined whether the different ultrastructural features correlated with the size of the synaptic active zone. We found that the volume, crista-membrane density and crista shape factor of presynaptic mitochondria are all strongly and significantly correlated with synapse size, (R=0.71, p=0.0005; R=0.77, p=0.0001; and R=0.65, p=0.0026, respectively; Pearson correlation, n=19 mitochondria from 2 mice, Fig. 3-1. B). We also tested our findings on post-mortem human tissue. We collected a random sample of mitochondria-containing presynaptic glutamatergic axon segments from the stratum radiatum of the hippocampal CA1 region, reconstructed them from serial sections (Fig. 3. H), and found a strong and significant correlation between the active zone area and the volume of individual presynaptic mitochondria (R=0.86, p<0.0001, n=31 mitochondria from 2 patients, Fig. 3. I, J), confirming the results from mouse studies. In summary, our results reveal for the first time that the ultrastructure of presynaptic mitochondria is coupled to synaptic performance in the brain, in a cell-type independent manner.

**Figure 3.**
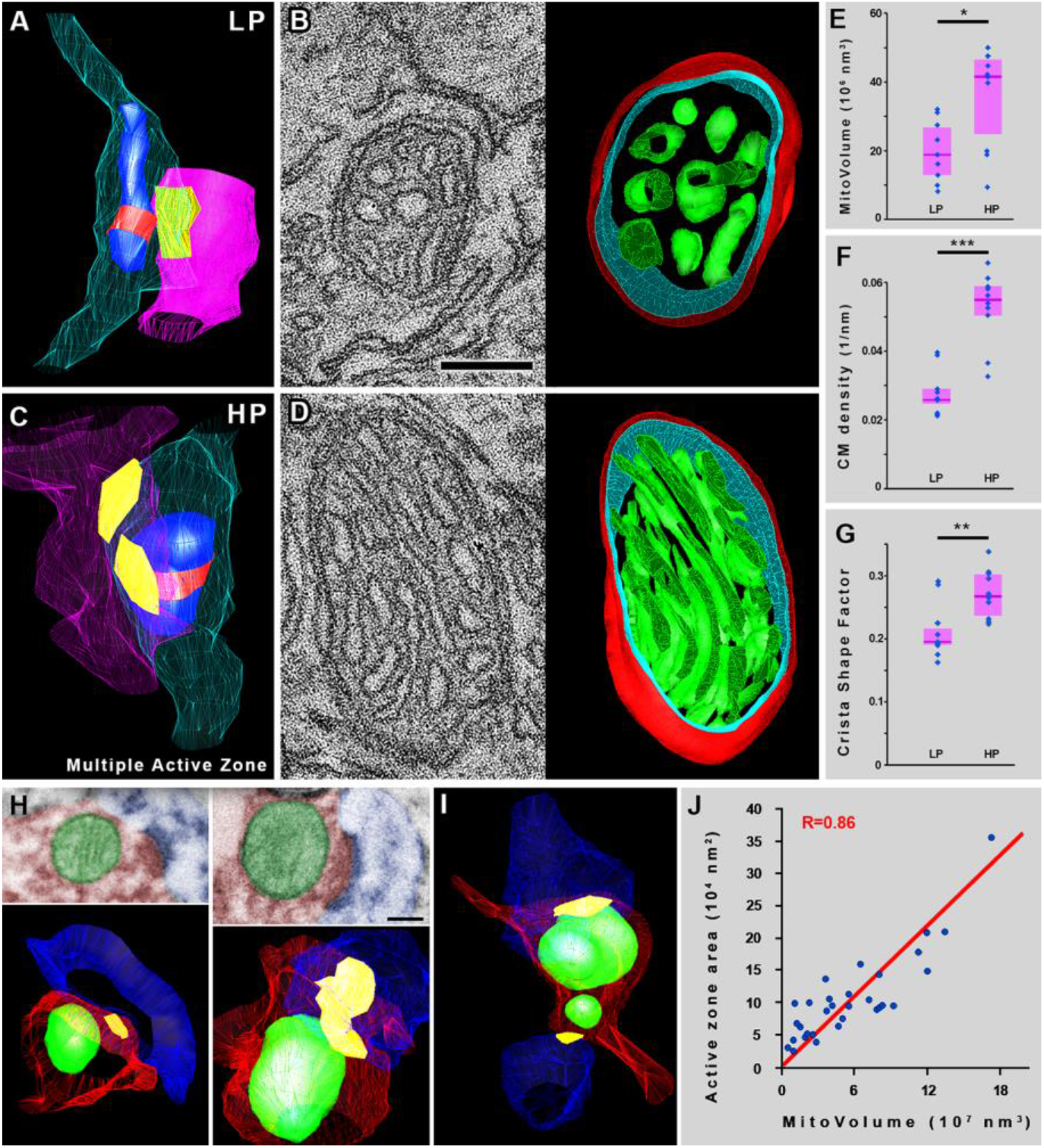
Mitochondrial ultrastructure is coupled to synaptic performance in a cell-type independent manner at glutamatergic synapses. (A) 3D-model of a serial EM reconstructed segment of a „low performance” glutamatergic bouton from the dentate-gyrus. Bouton membranes are semi-transparent cyan postsynaptic profiles are semi-transparent magenta, synapse are yellow, mitochondria are blue, red belts mark those sections of the organelles that were reconstructed through electron tomography. (B) 1.5 nm thick electron tomographic section and 3D-model of the reconstructed mitochondrion from the bouton depicted in A (mitochondrial outer membrane is red, inner boundary membrane is cyan, crista membrane is green). (C) 3D-model of a serial EM reconstructed segment of a „high performance” glutamatergic bouton from the dentate-gyrus (colors same as in A). (D) 1.5 nm thick electron tomographic section and 3D-model of the reconstructed mitochondrion from the bouton depicted in C. (E) Mitochondria are significantly larger in „high performance”-boutons than in „low performance”-boutons (Mann-Whitney U-test, p=0.0199, n=19 mitochondria from 2 mice). (F) Crista-membrane density is significantly higher in mitochondria of „high performance”-boutons than in those from „low performance”-boutons (Mann-Whitney U-test, p=0.0009, n=19 mitochondria from 2 mice). (G) Cristae are significantly more lamellar in mitochondria of „high performance”-boutons than in those from „low performance”-boutons, as verified by the higher crista shape factor values (Mann-Whitney U-test, p=0.0080, n=19 mitochondria from 2 mice). (H-J) The volume of individual presynaptic mitochondria correlates with active zone area in the human hippocampus. (H) Transmission electron micrographs of two presynaptic mitochondria from human samples, and their 3D-reconstructions from serial images. (I) 3D-model of an axonal segment from human tissue, giving two synapses, each with an associated presynaptic mitochondrion. Axons are red, mitochondria green, spines blue and active zones yellow. (J) The volume of presynaptic mitochondria shows a strong correlation with active zone area in human axons (R=0.86, p<0.0001, n=31 mitochondria from 2 patients). (E-G, J) Blue dots represent values from individual mitochondria, magenta rectangles represent interquartile ranges, deep-magenta lines mark median values. Scale bar is 110 nm in B and for D, 200 nm on H. The 3D models on B, D and H are not displayed on the same scale. See also Figures 3-1 and 3-2.

### Cytochrome-c density in presynaptic mitochondria of glutamatergic boutons scales with synaptic strength

To examine whether the observed cell-type independent coupling between mitochondrial ultrastructure and synaptic performance is accompanied also by changes in respiratory chain protein expression levels, we tested the density of CytC-labeling in mitochondria of „low-performance” (LP) and „high-performance” (HP) glutamatergic synaptic boutons, using correlated confocal and STORM super-resolution microscopy, taking advantage of the near-molecular resolution capability of this method. We labeled glutamategic boutons against vesicular glutamate transporter 1 (VG1), mitochondria with CytC, and postsynaptic densities with Homer1 (Fig. 4. A, C). We sampled mitochondria-containing synaptic boutons from the outer two-thirds of DG molecular layer, and reconstructed them from the deconvolved confocal stacks (Fig. 4. B, D). The volumes of Homer labeling belonging to each bouton were measured on the 3D models, and the median value was found to be 0.0265 μm^3^ (0.011-0.0518 interquartile range, Fig. 3-1. C). These values correspond to the size of the active zone in each bouton, meaning that this distribution indirectly reflects the output performance distribution of the examined boutons. Based on these data, we split the population at the median value, and sorted boutons with a lower Homer volume into the LP-group, and boutons with higher values into the HP-group (Fig. 3-1. C). We found that presynaptic mitochondria in HP-boutons were significantly larger than those in LP-boutons (HP-median: 0.0365 μm^3^, 0.0282-0.0578 interquartile range, LP-median: 0.0162 μm^3^, 0.0150-0.0265 interquartile range, 128.8% difference, Mann-Whitney U-test, p=0.0003, n=42 boutons from two mice; Fig. 4. G), and this observed difference is congruent with our serial EM data (128.8% vs. 121.6% difference). The CytC-labeling was recorded not only in the confocal channel, but also in the correlated STORM modality. The STORM SLP-s were overlayed the confocal images (Fig. 4. C, F).

**Figure 4.**
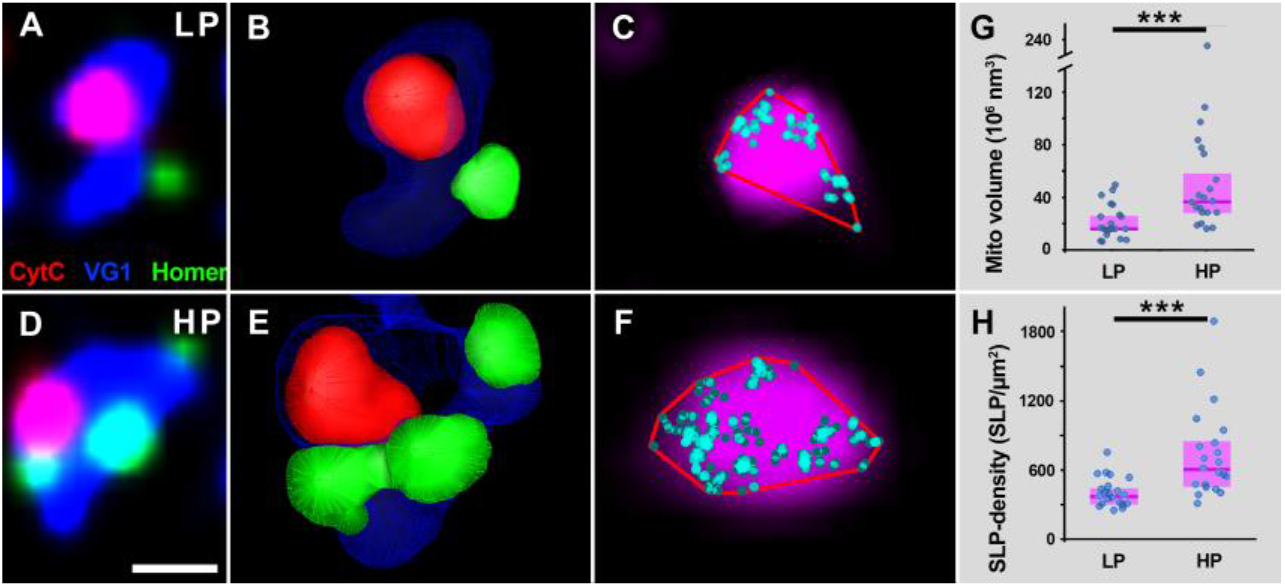
Cytochrome-c content of presynaptic mitochondria scales with synaptic performance at glutamatergic synapses (A) Confocal image shows a glutamatergic bouton (VG1, blue), containing a mitochondrion (Cyt-C, red), and its synapse (Homer1a, green). (B) The 3D-model of the bouton from A was reconstructed from the confocal stack. (C) Confocal and STORM image of mitochondrion enlarged from the same bouton (Cyt-C confocal in magenta, Cyt-C localisation points in cyan, 2D convex hull in red). (D-F) Another bouton depicted as in A-C. Mitochondria-containing synaptic boutons were divided into „low-performance” (LP) and „high-performance” (HP) groups, based on the corresponding Homer volumes. Bouton in A-C represents a LP, bouton in D-F represents a HP bouton. (G) Mitochondria in the HP-boutons are larger than those in the LP-boutons, as the CytC-volumes are significantly larger in the former group (Mann-Whitney U-test, p=0.00032, n=42 mitochondria from 2 mice). (H) Mitochondria in the HP-boutons contain cytochrome-c in a higher density than those in the HP-boutons (Mann-Whitney U-test, p=0.00016, n=42 mitochondria from 2 mice). Blue dots represent values from individual mitochondria, magenta rectangles represent interquartile ranges, deep-magenta lines mark median values. Scale bar is 800 nm for A and D, and 200 nm for C and F. The 3D models on B and E are not displayed on the same scale. See also Figure 3-1.

We found that the CytC SLP density was 61% higher in mitochondria of HP-boutons than in those of LP-boutons (HP-median: 598.5 SLP/ μm^2^, 455.6-822 interquartile range, LP-median: 370.8, 296.3-440.7 interquartile range, Mann-Whitney U-test, p=0.0001, n=42 boutons from two mice; Fig. 4. H). To verify the strength of our results, we pooled data from HP and LP boutons, and examined whether the investigated features correlated with the size of the synaptic active zone. We found, that the CytC-labeled volume and SLP-density are both strongly correlated to synapse size, (R=0.77, p<0.00001; R=0.60, p=0.00002, respectively; Spearman’s correlation, n=42 mitochondria from 2 mice, Fig. 3-1. D). Taken all these together, our results confirm that not only the mitochondrial ultrastructure is coupled to synaptic performance in a cell-type independent manner, but also the expression levels of the respiratory chain protein cytochrome-c.

## DISCUSSION

### Mitochondrial and synaptic performance are coupled via ultrastructure

The first observations suggesting a connection between mitochondrial crista-structure and function were obtained on *ex vivo* isolated mitochondria (Hackenbrock, 1966). Although those specific results have not been verified under physiological conditions, causal correlation between mitochondrial ultrastructure and output performance in non-neuronal cells has been confirmed in recent studies (Cogliati et al., 2016). The amount and density of cristae membrane as well as the lamellarity of cristae directly determine respiratory performance and efficiency (Cogliati et al., 2013; Else et al., 2004), and these ultrastructural features were also shown to influence performance at the level of the complete organism (Strohm and Daniels, 2003). On the other hand, aberrant morphology, dilatation or loss of mitochondrial cristae have been observed in a variety of pathological conditions (Choi et al., 2014; Daum et al., 2013), while genetic manipulations ameliorating damaged crista-structure have been shown to restore normal crista density/lamellarity and respiratory chain activity (Civiletto et al., 2015). Furthermore, modelling studies also confirmed, that crista-membrane density and lamellarity are key determinants of mitochondrial electrochemical potential and ATP-generating capacity (Song et al., 2013). There are several points explaining the correlation between mitochondrial ultrastructure and performance: larger crista-membrane surface can accommodate a higher amount of respiratory chain and ATP-synthase proteins, leading to a higher respiratory and ATP-generating capacity, whilst more lamellar cristae structure is beneficial for respiratory chain supercomplex assembly, resulting in a higher respiratory efficiency (Cogliati et al., 2013). Furthermore, more lamellar cristae – independently from changes in respiratory efficacy – result in an increased proton motive force, leading to higher ATP-producing capacity (Song et al., 2013). Finally, ATP-synthase molecules tend to be enriched and dimerize at curved cristae-rims, making the dense and tight packing of these membranes more effective to produce ATP (Davies et al., 2011).

Mitochondria are critically important for proper synaptic function, due to their central role in ATP-production, Ca^2+^-regulation and other major signalling mechanisms. In particular, presynaptic function has been shown to rely directly on activity-driven ATP synthesis (Rangaraju et al., 2014). The demand for mitochondrial function is reasonably coupled to neuronal activity (Gulyás et al., 2006; Kann et al., 2014) and also directly correlated with synaptic strength (Ivannikov et al., 2013; Smith et al., 2016; Sun et al., 2013; Verstreken et al., 2005). Our results demonstrate that increased synaptic performance is coupled to higher organelle volume, crista density, crista lamellarity and cytochrome-c levels at axonal release sites, suggesting that ultrastructure and molecular composition of presynaptic mitochondria are associated with synaptic performance.

### Performance coupling between mitochondria and synapses suggests a possible role for the activity dependent ultrastructural remodelling of mitochondria in neuroplasticity

Neuronal mitostasis – the maintenance of an appropriately distributed pool of healthy mitochondria – is fundamental for proper neuronal function (reviewed by Misgeld and Schwarz, 2017). Our observations verified the presence of a cell-type independent coupling between presynaptic mitochondrial ultrastructure and synaptic strength. This coupling – responsible for local demand matching – could be achieved either by a coordinated long-range retrograde/anterograde axonal trafficking of mitochondria, or by local plasticity mechanisms, that lead to adaptation at the level of single organelles.

During synaptic plasticity, the energetic demand of a potentiated synapse will increase. In case of the first mechanism – organelle trafficking – the mitochondrion from the potentiated bouton would need to be withdrawn, transported away – possibly back to the soma – and another, stronger mitochondrion would be sent out to replace it. The coordination of this process, together with the evident gap in energy supply during the change is more than problematic. On the other hand, the large number of papers dealing with mitochondrial motility and redistribution performed mainly *in vitro* experiments, however, recent *in vivo* two-photon imaging studies suggest that mitochondrial motility is way much smaller, and also quite independent from neuronal activity (Lewis et al., 2016; Smit-Rigter et al., 2016). This suggests that this mechanism alone is not sufficient to account for local demand matching. The other possible mechanism is locally regulated ultrastructural plasticity. In fact, in non-neuronal tissues, an overwhelming body of evidence confirms that dynamic ultrastructural remodeling of mitochondria takes place as a response to altered energetic demand (Cogliati et al., 2016).

Mitochondrial inner membrane architecture has been shown to undergo substantial remodelling after exercise, under hypoxic conditions or during starvation (Gomes et al., 2011; Hambrecht et al., 1997; Nielsen et al., 2016; Perkins et al., 2012), suggesting that the local demand for mitochondrial performance is communicated to these organelles, prompting their activity-dependent ultrastructural plasticity. The common response to an increased energetic demand was an increased crista-membrane density and lamellarity in all of these cases. The idea of mitochondrial ultrastructural adaptation is also supported by a study, confirming that crista remodelling can happen on the timescale of minutes (Dikov and Bereiter-Hahn, 2013). In recent years, various mechanisms have been described to regulate mitochondrial ultrastructure, such as mitochondrial dynamin like GTPase (OPA1), mitochondrial contact site and cristae organizing system (MICOS), ATP-synthase dimerisation and the inner-membrane protein Pantagruelian Mitochondrion I (Gomes et al., 2011; Hahn et al., 2016; Harner et al., 2011; Macchi et al., 2013; Neupert, 2012). Substrate dependent changes in OPA1 oligomer levels (Patten et al., 2014; Sood et al., 2014), or activity-dependent ATP-synthase clustering (Jimenez et al., 2014) are known to contribute to mitochondrial ultrastructural plasticity. Synaptic mitochondria possess a specific proteome (Völgyi et al., 2015), and local axonal protein synthesis has also been shown to be required for the maintenance of mitochondrial function (Aschrafi et al., 2008; Gale et al., 2017; Hillefors et al., 2007). Thus, it is very likely that local presynaptic signalling mechanisms regulate the expression of mitochondrial proteins in axons, thereby controlling mitochondrial ultrastructure and performance.

All these findings – together with our results – suggest that the primary way of adjusting mitochondrial performance to the actual demand at synapses could be the activity-dependent ultrastructural plasticity of these organelles. Furthermore, our results – confirming a cell-type independent coupling between synaptic performance and mitochondrial ultrastructure – indicate that these changes in mitochondrial ultrastructure and molecular fingerprints could contribute to neuroplasticity at the level of individual synapses.

## AUTHOR CONTRIBUTIONS

Methodology C.C.; Investigation C.C. B. P. and A.S.; Conceptualization and Writing – Original Draft, C.C.; Writing – Review & Editing, C.C. and A.D.; Funding Acquisition, A.D.; Resources, C.C. and A.D.; Supervision, C.C. and A.D.

## ACKNOWLEDGEMENTS

This work was supported by “Momentum” research grant from the Hungarian Academy of Sciences (LP2016-4/2016 to A.D.) and ERC-CoG 724994 (A.D.). We thank László Barna and the Nikon Imaging Center at the Institute of Experimental Medicine for kindly providing microscopy support, and David Mastronarde at MCDB for his continuous help with IMOD software. We thank the Department of Pathology, St. Borbála Hospital, Tatabánya, and the Human Brain Research Lab at the Institute of Experimental Medicine, Hungarian Academy of Sciences (IEM HAS) for providing human brain tissue. We also thank Dr. Gábor Nyiri, Prof. Ken Mackie and Prof. Andreas Zimmer for the CB1R antibodies and CB1R-KO animals; and Dóra Gali-Györkei for her excellent technical assistance.

The Authors report no conflict of interest.

## EXTENDED DATA

**Figure 1-1.**
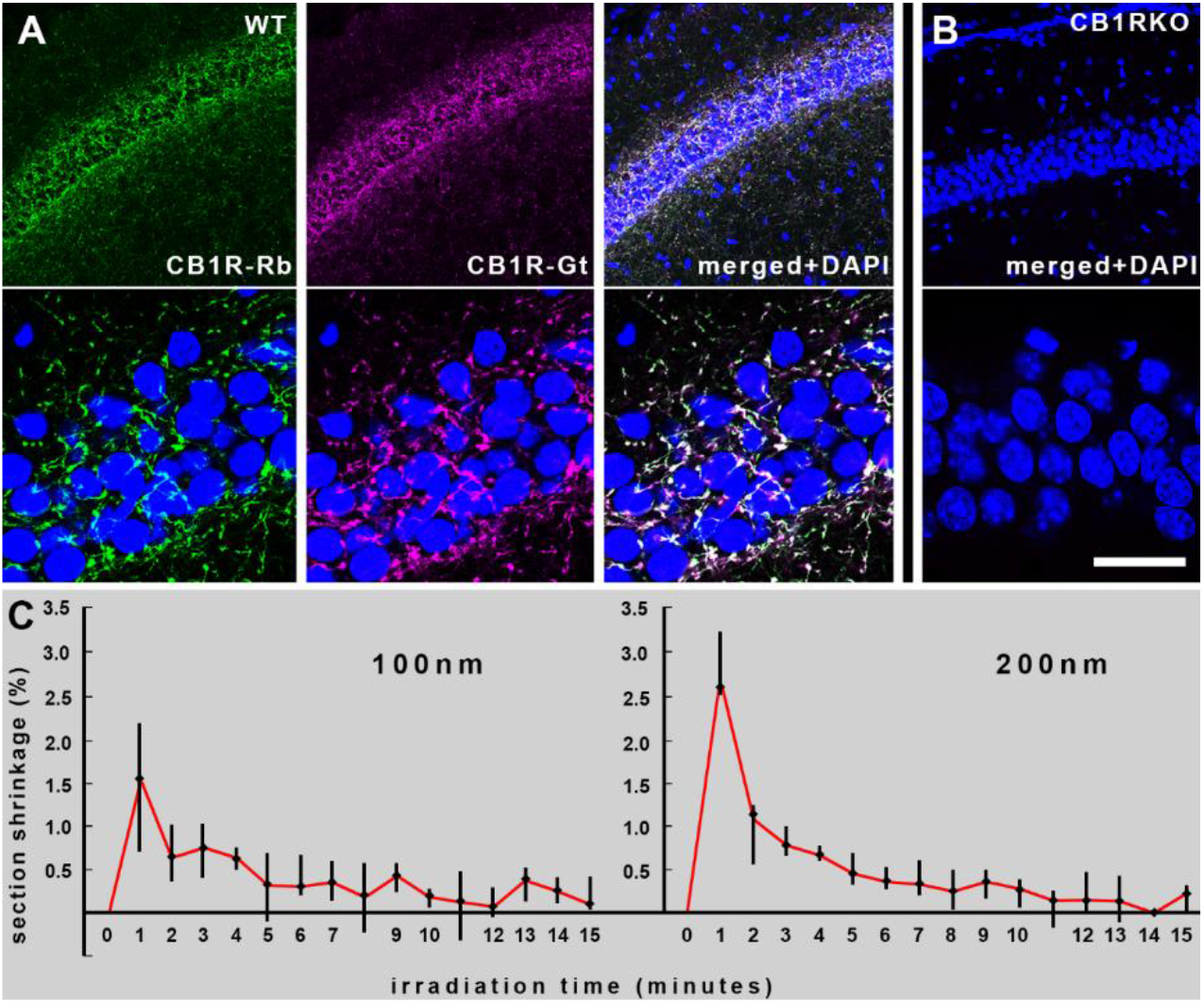
Staining with two anti-CB1R antibodies is overlapping in WT, but totally absent from CB1RKO mice; rate of tissue shrinkage due to electron beam irradiation. **A** Triple-color confocal images show CB1R staining in the hippocampal CA1 region of a wild-type mouse with a rabbit (green) and a goat (magenta) antibody. Cell nuclei are stained with DAPI (blue). **B** Staining with both antibodies are completely absent in CB1R-KO mice. Scale bar is 100 μm for the upper and 30 μm for the lower panels. **C** Rates of tissue shrinkage on 100 (left) and 200 nm thick sections (right) under 15 minutes of electron beam irradiation. Black bars represent interquartile ranges, black dots mark median values.

**Figure 1-2.**
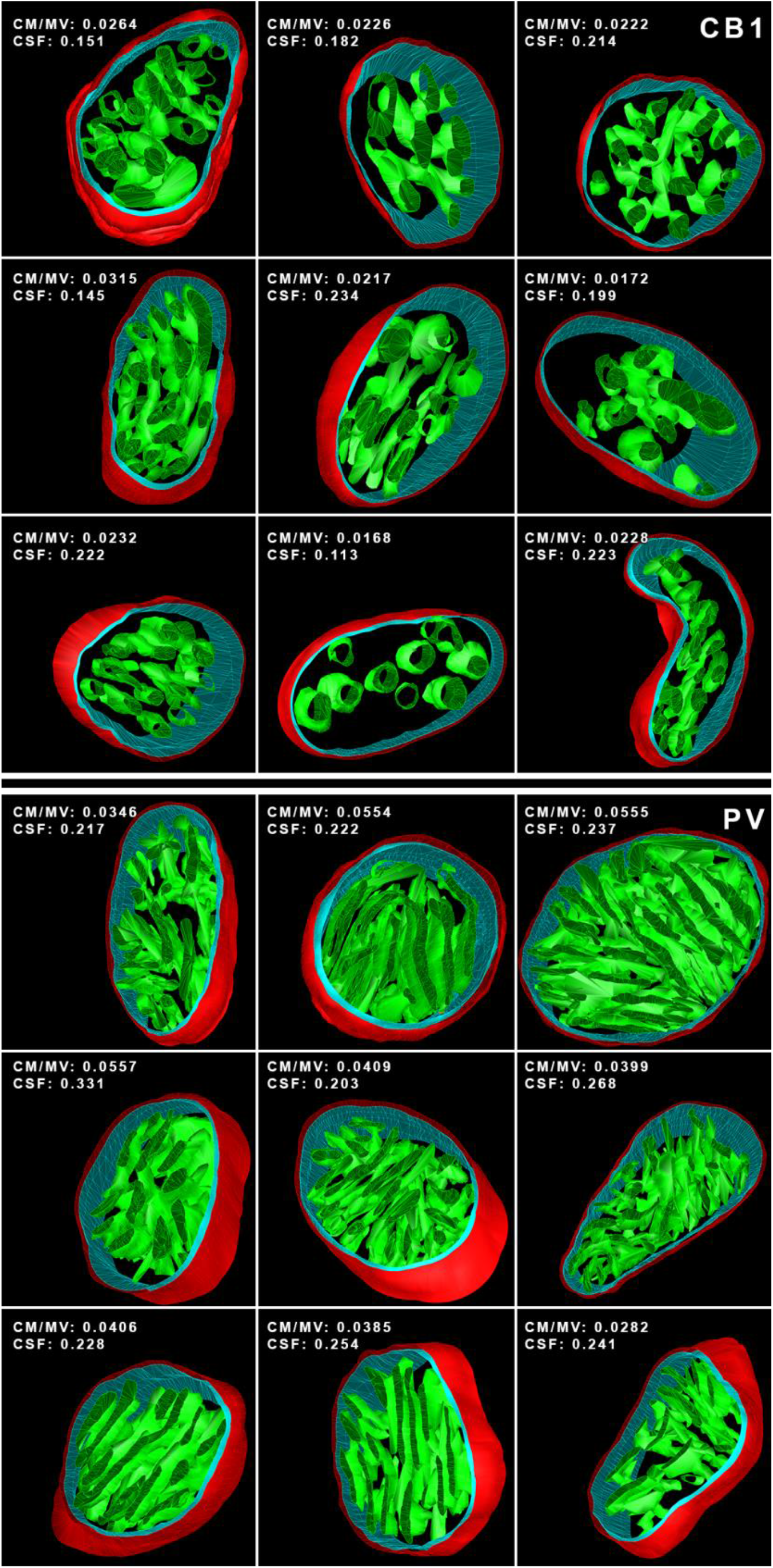
3D models of the reconstructed mitochondrial volumes from the hippocampal CA1 region. Top three rows show presynaptic mitochondria from regular spiking basket cell boutons, while the bottom three rows from fast-spiking basket-cell boutons. Mitochondrial outer membrane is red, inner boundary membrane is turquise, crista membrane is green. CM/MV: crista membrane area/mitochondrial volume, CSF: crista shape factor. The models are not displayed on the same magnification.

**Figure 3-1.**
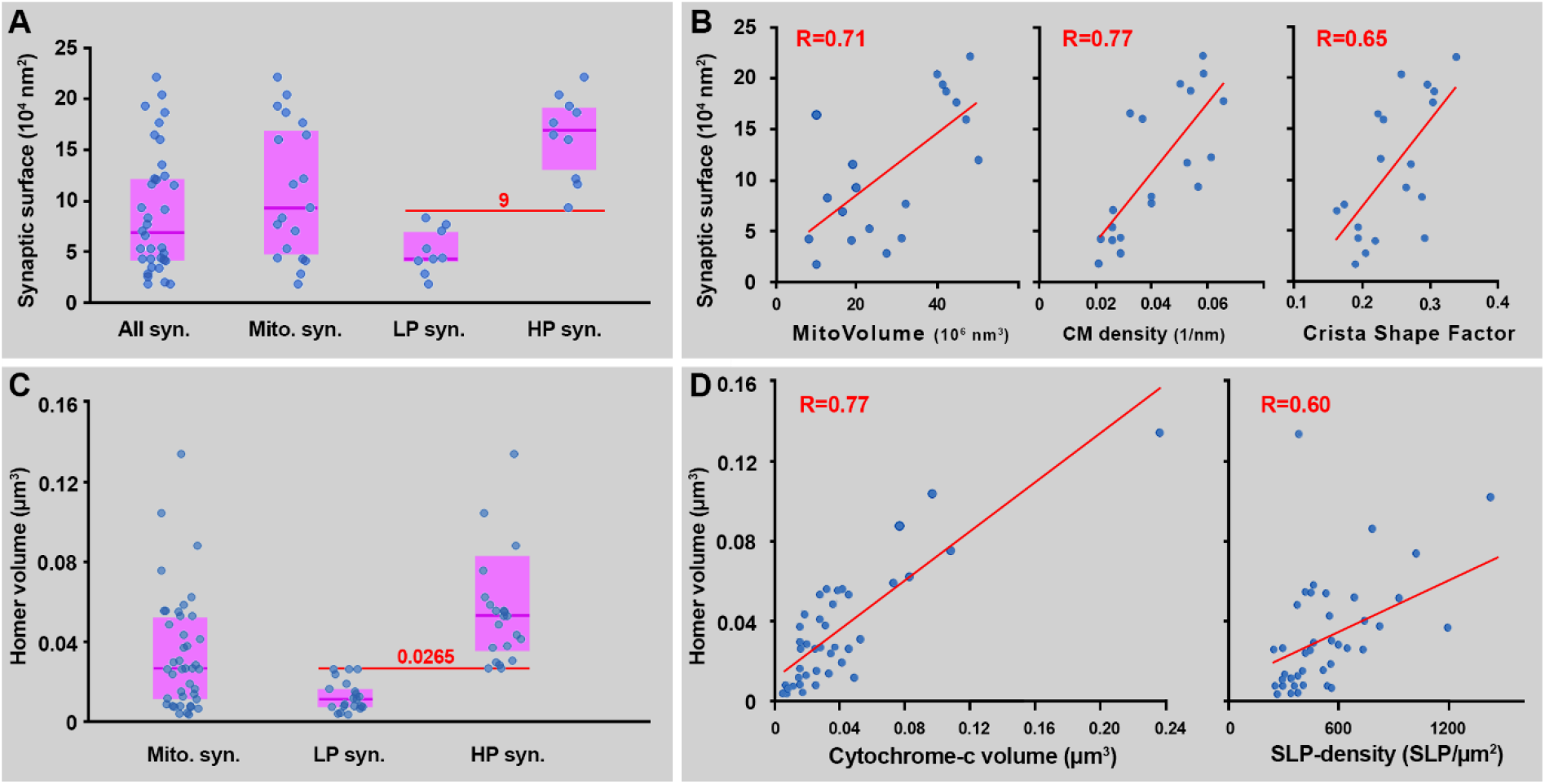
Distribution of active zone sizes of the glutamatergic boutons in the tomographic and STORM measurements, and correlations of performance-determining mitochondrial features with synapse size. **A** Active zone area distribution of the glutamatergic boutons examined with serial electron microscopy and electron tomography. **B** The volume, crista-membrane density and crista shape factor of presynaptic mitochondria are all strongly correlated to synapse size, (R=0.71, p=0.0005; R=0.77, p=0.0001; and R=0.65, p=0.0026, respectively). Blue dots represent values from individual mitochondria, regression lines are red. **C** Active zone area distribution of the glutamatergic boutons examined with confocal laser scanning microscopy and STORM super-resolution microscopy. (All syn.: all examined synaptic boutons, Mito. syn.: synaptic boutons with mitochondria, LP syn.: low performance synaptic boutons, HP syn.: high performance synaptic boutons). Blue dots represent values from individual boutons, magenta rectangles represent interquartile ranges, deep-magenta lines mark median values. Red lines show values dividing HP and LP synaptic boutons. **D** The cytochrome-c labeled volume and SLP-density are both strongly correlated to synapse size, (R=0.77, p<0.00001; R=0.60, p=0.00002, respectively). Blue dots represent values from individual mitochondria, regression lines are red.

**Figure 3-2.**
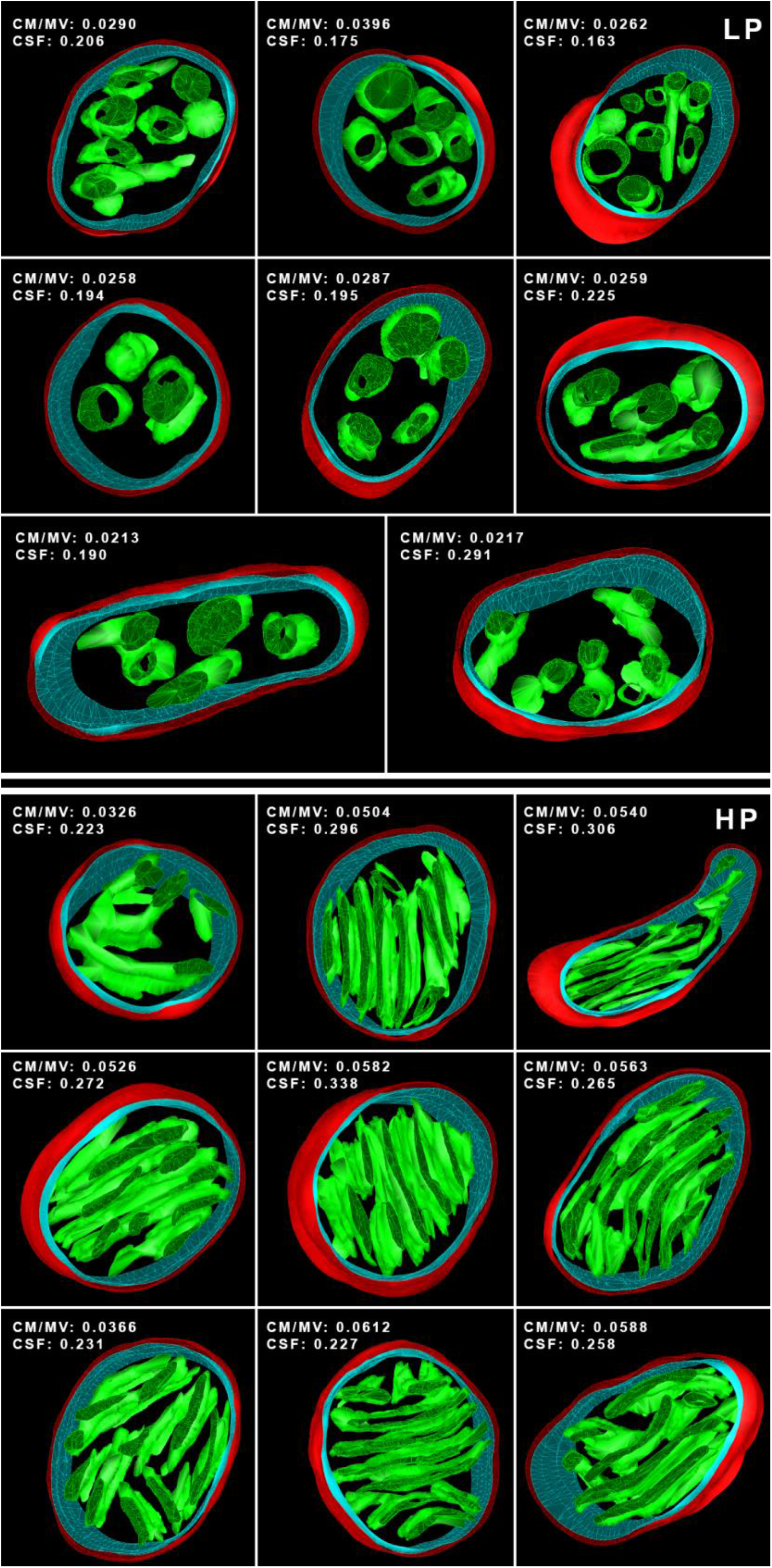
3D models of the reconstructed mitochondrial volumes from the dentate gyrus. Top three rows show presynaptic mitochondria from “low performance” glutamatergic boutons, while the bottom three rows from „high performance” ones. Mitochondrial outer membrane is red, inner boundary membrane is turquise, crista membrane is green. CM/MV: crista membrane area/mitochondrial volume, CSF: crista shape factor. The models are not displayed on the same magnification.

## REFERENCES

Amaral, D.G., Scharfman, H.E., and Lavenex, P. (2007). The dentate gyrus: fundamental neuroanatomical organization (dentate gyrus for dummies). Prog. Brain Res.

Andreska, T., Aufmkolk, S., Sauer, M., and Blum, R. (2014). High abundance of BDNF within glutamatergic presynapses of cultured hippocampal neurons. Front. Cell. Neurosci. 8, 1–15.

Aschrafi, A., Schwechter, A.D., Mameza, M.G., Natera-Naranjo, O., Gioio, A.E., and Kaplan, B.B. (2008). MicroRNA-338 regulates local cytochrome c oxidase IV mRNA levels and oxidative phosphorylation in the axons of sympathetic neurons. J. Neurosci. 28, 12581–12590.

Barna, L., Dudok, B., Miczán, V., Horváth, A., László, Z.I., and Katona, I. (2016). Correlated confocal and super-resolution imaging by VividSTORM. Nat. Protoc. 11, 163–183.

Barnhart, E.L. (2016). Mechanics of mitochondrial motility in neurons. Curr. Opin. Cell Biol.

Billups, B., and Forsythe, I.D. (2002). Presynaptic mitochondrial calcium sequestration influences transmission at mammalian central synapses. J. Neurosci. 22, 5840–5847.

Bourne, J.N., and Harris, K.M. (2012). Nanoscale analysis of structural synaptic plasticity. Curr. Opin. Neurobiol. 22, 372–382.

Buchs, P.A., and Muller, D. (1996). Induction of long-term potentiation is associated with major ultrastructural changes of activated synapses. Proc. Natl. Acad. Sci. U. S. A. 93, 8040–8045.

Chandel, N.S. (2014). Mitochondria as signaling organelles. BMC Biol. 12.

Cheetham, C.E.J., Barnes, S.J., Albieri, G., Knott, G.W., and Finnerty, G.T. (2014). Pansynaptic enlargement at adult cortical connections strengthened by experience. Cereb. Cortex 24, 521–531.

Choi, K.J., Kim, M.J., Je, A.R., Jun, S., Lee, C., Lee, E., Jo, M., Huh, Y.H., and Kweon, H.S. (2014). Three-dimensional analysis of abnormal ultrastructural alteration in mitochondria of hippocampus of APP/PSEN1 transgenic mouse. J. Biosci.

Civiletto, G., Varanita, T., Cerutti, R., Gorletta, T., Barbaro, S., Marchet, S., Lamperti, C., Viscomi, C., Scorrano, L., and Zeviani, M. (2015). Opa1 overexpression ameliorates the phenotype of two mitochondrial disease mouse models. Cell Metab.

Cogliati, S., Frezza, C., Soriano, M.E., Varanita, T., Quintana-Cabrera, R., Corrado, M., Cipolat, S., Costa, V., Casarin, A., Gomes, L.C., et al. (2013). Mitochondrial cristae shape determines respiratory chain supercomplexes assembly and respiratory efficiency. Cell.

Cogliati, S., Enriquez, J.A., and Scorrano, L. (2016). Mitochondrial Cristae: Where Beauty Meets Functionality. Trends Biochem. Sci.

Daum, B., Walter, A., Horst, A., Osiewacz, H.D., and Kühlbrandt, W. (2013). Age-dependent dissociation of ATP synthase dimers and loss of inner-membrane cristae in mitochondria. Proc. Natl. Acad. Sci. U. S. A. 110, 15301–15306.

Davies, K.M., Strauss, M., Daum, B., Kief, J.H., Osiewacz, H.D., Rycovska, A., Zickermann, V., and Kuhlbrandt, W. (2011). Macromolecular organization of ATP synthase and complex I in whole mitochondria. Proc. Natl. Acad. Sci. 108, 14121–14126.

Desmond, N.L., and Levy, W.B. (1988). Synaptic interface surface area increases with long-term potentiation in the hippocampal dentate gyrus. Brain Res 453, 308–314.

Dikov, D., and Bereiter-Hahn, J. (2013). Inner membrane dynamics in mitochondria. J. Struct. Biol.

Else, P.L., Turner, N., and Hulbert, A.J. (2004). The evolution of endothermy: role for membranes and molecular activity. Physiol. Biochem. Zool. 77, 950–958.

Gale, J.R., Aschrafi, A., Gioio, A.E., and Kaplan, B.B. (2017). Nuclear-Encoded Mitochondrial mRNAs: A Powerful Force in Axonal Growth and Development. Neurosci. 107385841771422.

Gazit, N., Vertkin, I., Shapira, I., Helm, M., Slomowitz, E., Sheiba, M., Mor, Y., Rizzoli, S., and Slutsky, I. (2016). IGF-1 Receptor Differentially Regulates Spontaneous and Evoked Transmission via Mitochondria at Hippocampal Synapses. Neuron.

Geinisman, Y. (2000). Structural synaptic modifications associated with hippocampal LTP and behavioral learning. Cereb. Cortex 10, 952–962.

Gomes, L.C., Di Benedetto, G., and Scorrano, L. (2011). During autophagy mitochondria elongate, are spared from degradation and sustain cell viability. Nat. Publ. Gr. 13.

Gulyás, A.I., Buzsáki, G., Freund, T.F., and Hirase, H. (2006). Populations of hippocampal inhibitory neurons express different levels of cytochrome c. Eur. J. Neurosci.

Gunter, T.E., Buntinas, L., Sparagna, G., Eliseev, R., and Gunter, K. (2000). Mitochondrial calcium transport: mechanisms and functions. Cell Calcium 28, 285–296.

Hackenbrock, C.R. (1966). Ultrastructural bases for metabolically linked mechanical activity in mitochondria. I. Reversible ultrastructural changes with change in metabolic steady state in isolated liver mitochondria. J. Cell Biol. 30, 269–297.

Hahn, A., Parey, K., Bublitz, M., Mills, D.J., Zickermann, V., Vonck, J., Kühlbrandt, W., and Meier, T. (2016). Structure of a Complete ATP Synthase Dimer Reveals the Molecular Basis of Inner Mitochondrial Membrane Morphology. Mol. Cell.

Hall, C.N., Klein-Flugge, M.C., Howarth, C., and Attwell, D. (2012). Oxidative Phosphorylation, Not Glycolysis, Powers Presynaptic and Postsynaptic Mechanisms Underlying Brain Information Processing. J. Neurosci. 32, 8940–8951.

Hambrecht, R., Fiehn, E., Yu, J., Niebauer, J., Weigl, C., Hilbrich, L., Adams, V., Riede, U., and Schuler, G. (1997). Effects of endurance training on mitochondrial ultrastructure and fiber type distribution in skeletal muscle of patients with stable chronic heart failure. J. Am. Coll. Cardiol. 29, 1067–1073.

Harner, M., Körner, C., Walther, D., Mokranjac, D., Kaesmacher, J., Welsch, U., Griffith, J., Mann, M., Reggiori, F., and Neupert, W. (2011). The mitochondrial contact site complex, a determinant of mitochondrial architecture. EMBO J. 30, 4356–4370.

Harris, J.J., Jolivet, R., and Attwell, D. (2012). Synaptic Energy Use and Supply. Neuron.

Hillefors, M., Gioio, A.E., Mameza, M.G., and Kaplan, B.B. (2007). Axon viability and mitochondrial function are dependent on local protein synthesis in sympathetic neurons. Cell. Mol. Neurobiol.

Holderith, N., Lorincz, A., Katona, G., Kulik, A., Watanabe, M., and Nusser, Z. (2012). Release probability of hippocampal glutamatergic terminals scales with the size of the active zone. Nat. Neurosci.

Holtmaat, A., and Svoboda, K. (2009). Experience-dependent structural synaptic plasticity in the mammalian brain. Nat. Rev. Neurosci. 10, 759–759.

Huang, B., Jones, S.A., Brandenburg, B., and Zhuang, X. (2008). Whole-cell 3D STORM reveals interactions between cellular structures with nanometer-scale resolution. Nat. Methods 5, 1047–1052.

Ivannikov, M. V., Sugimori, M., and Llinás, R.R. (2013). Synaptic vesicle exocytosis in hippocampal synaptosomes correlates directly with total mitochondrial volume. J. Mol. Neurosci.

Jimenez, L., Laporte, D., Duvezin-Caubet, S., Courtout, F., and Sagot, I. (2014). Mitochondrial ATP synthases cluster as discrete domains that reorganize with the cellular demand for oxidative phosphorylation. J. Cell Sci. 127, 719–726.

Kann, O., Papageorgiou, I.E., and Draguhn, A. (2014). Highly energized inhibitory interneurons are a central element for information processing in cortical networks. J. Cereb. Blood Flow Metab. 34, 1270–1282.

Klausberger, T., Marton, L.F., O’Neill, J., Huck, J.H.J., Dalezios, Y., Fuentealba, P., Suen, W.Y., Papp, E., Kaneko, T., Watanabe, M., et al. (2005). Complementary roles of cholecystokinin- and parvalbumin-expressing GABAergic neurons in hippocampal network oscillations. J. Neurosci.

Korogod, N., Petersen, C.C.H., and Knott, G.W. (2015). Ultrastructural analysis of adult mouse neocortex comparing aldehyde perfusion with cryo fixation. Elife.

Kremer, J.R., Mastronarde, D.N., and Mcintosh, J.R. (1996). Computer Visualization of ThreeDimensional Image Data Using IMOD. J. Struct. Biol. 116, 71–76.

Lapray, D., Lasztoczi, B., Lagler, M., James Viney, T., Katona, L., Valenti, O., Hartwich, K., Borhegyi, Z., Somogyi, P., and Klausberger, T. (2012). Behavior-dependent specialization of identified hippocampal interneurons. Nat. Neurosci. 15.

Lewis, T.L., Turi, G.F., Kwon, S.K., Losonczy, A., and Polleux, F. (2016). Progressive Decrease of Mitochondrial Motility during Maturation of Cortical Axons In Vitro and In Vivo. Curr. Biol. 26, 2602–2608.

Li, Z., Okamoto, K.I., Hayashi, Y., and Sheng, M. (2004). The importance of dendritic mitochondria in the morphogenesis and plasticity of spines and synapses. Cell.

Macchi, M., El Fissi, N., Tufi, R., Bentobji, M., Liévens, J.-C., Martins, L.M., Royet, J., and Rival, T. (2013). The Drosophila inner-membrane protein PMI controls crista biogenesis and mitochondrial diameter. J. Cell Sci. 126, 814–824.

Mannella, C.A., Lederer, W.J., and Jafri, M.S. (2013). The connection between inner membrane topology and mitochondrial function. J Mol Cell Cardiol 62C, 51–57.

Melone, M., Burette, A., and Weinberg, R.J. (2005). Light microscopic identification and immunocytochemical characterization of glutamatergic synapses in brain sections. J. Comp. Neurol.

Misgeld, T., and Schwarz, T.L. (2017). Mitostasis in Neurons: Maintaining Mitochondria in an Extended Cellular Architecture. Neuron 96, 651–666.

Muller, D., Nikonenko, I., Jourdain, P., and Alberi, S. (2002). LTP, memory and structural plasticity. Curr. Mol. Med. 2, 605–611.

Neupert, W. (2012). SnapShot: Mitochondrial Architecture. Cell 149, 722–722.e1.

Nicastro, D., Frangakis, A.S., Typke, D., and Baumeister, W. (2000). Cryo-electron tomography of neurospora mitochondria. J. Struct. Biol. 129, 48–56.

Nielsen, J., Gejl, K.D., Hey-mogensen, M., Holmberg, H.-C., Suetta, C., Krustrup, P., Elemans, C.P.H., and Ørtenblad, N. (2016). Plasticity in mitochondrial cristae density allows metabolic capacity modulation in human skeletal muscle Corresponding author: Key points. J. Physiol. 1–29.

Patten, D.A., Wong, J., Khacho, M., Soubannier, V., Mailloux, R.J., Pilon-larose, K., Maclaurin, J.G., Park, D.S., Mcbride, H.M., Trinkle-mulcahy, L., et al. (2014). OPA 1 -dependent cristae modulation is essential for cellular adaptation to metabolic demand. 33, 2676–2691.

Perkins, G., Hsiao, Y. hsin, Yin, S., Tjong, J., Tran, M.T., Lau, J., Xue, J., Liu, S., Ellisman, M.H., and Zhou, D. (2012). Ultrastructural Modifications in the Mitochondria of Hypoxia-Adapted Drosophila melanogaster. PLoS One.

Perkins, G.A., Song, J.Y., Tarsa, L., Deerinck, T.J., Ellisman, M.H., and Frey, T.G. (1998). Electron tomography of mitochondria from brown adipocytes reveals crista junctions. J. Bioenerg. Biomembr.

Perkins, G. a, Ellisman, M.H., and Fox, D. a (2003). Three-dimensional analysis of mouse rod and cone mitochondrial cristae architecture: bioenergetic and functional implications. Mol. Vis. 9, 60–73.

Popov, V.I., Davies, H.A., Rogachevsky, V. V., Patrushev, I. V., Errington, M.L., Gabbott, P.L.A., Bliss, T.V.P., and Stewart, M.G. (2004). Remodelling of synaptic morphology but unchanged synaptic density during late phase long-term potentiation (LTP): A serial section electron micrograph study in the dentate gyrus in the anaesthetised rat. Neuroscience.

Rangaraju, V., Calloway, N., and Ryan, T.A. (2014). Activity-driven local ATP synthesis is required for synaptic function. Cell 156, 825–835.

Sheng, Z.H. (2014). Mitochondrial trafficking and anchoring in neurons: New insight and implications. J. Cell Biol. 204, 1087–1098.

Smit-Rigter, L., Rajendran, R., Silva, C.A.P., Spierenburg, L., Groeneweg, F., Ruimschotel, E.M., van Versendaal, D., van der Togt, C., Eysel, U.T., Heimel, J.A., et al. (2016). Mitochondrial Dynamics in Visual Cortex Are Limited In Vivo and Not Affected by Axonal Structural Plasticity. Curr. Biol. 26, 2609–2616.

Smith, H.L., Bourne, J.N., Cao, G., Chirillo, M.A., Ostroff, L.E., Watson, D.J., and Harris, K.M. (2016). Mitochondrial support of persistent presynaptic vesicle mobilization with age-dependent synaptic growth after LTP. Elife 5, e15275.

Song, D.H., Park, J., Maurer, L.L., Lu, W., Philbert, M.A., and Sastry, A.M. (2013). Biophysical significance of the inner mitochondrial membrane structure on the electrochemical potential of mitochondria. Phys. Rev. E. Stat. Nonlin. Soft Matter Phys.

Sood, A., Jeyaraju, D.V., Prudent, J., Caron, A., Lemieux, P., McBride, H.M., Laplante, M., Toth, K., and Pellegrini, L. (2014). A Mitofusin-2-dependent inactivating cleavage of Opa1 links changes in mitochondria cristae and ER contacts in the postprandial liver. Proc. Natl. Acad. Sci. U. S. A. 111, 16017–16022.

Strohm, E., and Daniels, W. (2003). Ultrastructure meets reproductive success: performance of a sphecid wasp is correlated with the fine structure of the flight-muscle mitochondria.

Sun, T., Qiao, H., Pan, P.Y., Chen, Y., and Sheng, Z.H. (2013). Motile axonal mitochondria contribute to the variability of presynaptic strength. Cell Rep.

Takács, V.T., Szőnyi, A., Freund, T.F., Nyiri, G., and Gulyás, A.I. (2015). Quantitative ultrastructural analysis of basket and axo-axonic cell terminals in the mouse hippocampus. Brain Struct. Funct.

Toussaint, C., and Kugler, P. (1989). Anatomy and Embryology Morphometric analysis of mitochondria and boutons in the dentate gyrus molecular layer of aged rats. Anat Embryol 179, 411–414.

Verstreken, P., Ly, C. V., Venken, K.J.T., Koh, T.W., Zhou, Y., and Bellen, H.J. (2005). Synaptic mitochondria are critical for mobilization of reserve pool vesicles at Drosophila neuromuscular junctions. Neuron.

Völgyi, K., Gulyássy, P., Háden, K., Kis, V., Badics, K., Kékesi, K.A., Simor, A., Györffy, B., Tóth, E. A., Lubec, G., et al. (2015). Synaptic mitochondria: A brain mitochondria cluster with a specific proteome. J. Proteomics.

Wilson, D.F., Harrison, D.K., and Vinogradov, A. (2014). Mitochondrial cytochrome c oxidase and control of energy metabolism: measurements in suspensions of isolated mitochondria. J. Appl. Physiol. 117, 1424–1430.

Zimmer, A., Zimmer, A.M., Hohmann, A.G., Herkenham, M., and Bonner, T.I. (1999). Increased mortality, hypoactivity, and hypoalgesia in cannabinoid CB1 receptor knockout mice. Neurobiology 96, 5780–5785.

